# Whole genome sequencing of endometrial cancer identifies novel subgroups, drivers, and actionable alterations

**DOI:** 10.64898/2026.06.15.730391

**Authors:** Sebastian Meyer, Ben Kinnersley, Katarzyna Kedzierska, Ignacio Soriano, Eszter Lakatos, Claudia Arnedo-Pac, Richard Culliford, Avraam Tapinos, Laura Knight, Yannick Comoglio, Anna Frangou, Alex J. Cornish, Aliah Hawari, Daniel Chubb, Amit Sud, Boris Noyvert, Steve Thorn, Helen White, Alona Sosinsky, Ahmed Ahmed, James D Brenton, Nuria Lopez-Bigas, Andrea Sottoriva, Tjalling Bosse, Emma J. Davidson, 100kGP Endometrial Cancer Clinical Consortium, Richard Edmondson, Trevor Graham, Ian Tomlinson, Richard S. Houlston, Andreas J Gruber, David C. Wedge, David N Church

## Abstract

Endometrial cancer (EC) is the most common gynaecological malignancy in high income countries, and is increasing in incidence. While molecular stratification has improved its management, precision care is hampered by incomplete characterization of the EC genome. We address this by analysis of whole genome sequencing (WGS) of 665 ECs generated by the UK Genomics England 100,000 Genome Project (100kGP). 5% of cases were associated with germline pathogenic variants in cancer genes, including *BRCA1* which we confirmed predisposes to EC. We identified 107 putative coding driver genes, 35% of which had no prior established role in EC. Novel structural variants included gains of *MYCN* and loss of its negative regulator NEDD4.1 which were significantly mutually exclusive in copy number (CN) high tumours. Immunogenomic analysis confirmed selection for driver alterations of low immunogenicity based on patient HLA haplotype, and pervasive immune escape through multiple mechanisms. Unsupervised clustering of mutational signatures and genomic alterations identified known and novel molecular subgroups, including a CN-high subset with mutational signatures of homologous recombination deficiency (HRD) and favourable outcome. Independent prognostic value of single nucleotide variant (SNV) burden, CN burden and multiple coding drivers, along with the identification of targetable molecular alterations in over one-third of cases, underscores the promise of WGS for precision medicine in EC.

## INTRODUCTION

Endometrial cancer (EC) is the commonest gynaecological malignancy in high income countries^1^, and is increasing in incidence^2^. Following a landmark study of 248 EC exomes by The Cancer Genome Atlas^3^, the traditional classification of EC by histology has been supplanted by a molecular classifier comprising four subgroups which vary in single nucleotide variant (SNV) and copy number (CN) alteration burden. These comprise: (i) tumours with ultra-high SNV burden caused by somatic DNA polymerase epsilon (*POLE*) exonuclease domain mutations; (ii) tumours with microsatellite instability (MSI), elevated SNV and insertion-deletion (indel) mutation burden (henceforth collectively referred to as small somatic variants – SSVs – unless individually specified) caused by DNA mismatch repair deficiency (MMRd); (iii) tumours with low SSV burden and few copy number (CN) alterations (CN-low); and (iv) SSV low tumours with high CN alteration burden (CN-high). TCGA and other exome sequencing studies have identified alterations in genes, pathways and processes critical for EC development, including recurrent alterations in the PI3K and MAPK pathways, mechanisms of genomic instability, and dysregulation of chromatin organization^4–6^. However, they have limitations. None have analysed the whole EC and corresponding normal genome to comprehensively identify germline predisposition, mutational signatures, coding, non-coding and structural drivers and the relationship between these factors. Nor have they systematically analysed the potential of these alterations for risk stratification or therapeutic targeting, both of which remain largely unknown. As genomic sequencing becomes incorporated into clinical management of EC, this represents an increasing unmet need. Here, we address this by analysing whole genome sequencing (WGS) of 665 ECs generated by the UK Genomics England 100,000 Genome Project (100kGP)^7^. We identify novel tumour subgroups, drivers, clinically actionable alterations, and provide a roadmap for the implementation of WGS for EC in clinical practice.

## RESULTS

### Patient and tumour characteristics

We analysed WGS data from 665 ECs and paired normal samples (median sequencing depth 100x and 30x respectively) from patients undergoing routine clinical care in the UK. Cases were unselected for patient or tumour characteristics (median age 68, range 26–96). The vast majority of cases (n=657; 98.8%) were localized primary tumours resected by hysterectomy, and most (71%) were of endometrioid histotype, though serous (14%), carcinosarcomas (7.2%) and other uncommon histologies were included at expected frequencies. Following comprehensive clinical and molecular quality control, tumours were classified into molecular EC subgroups^4^ based on MSI status (determined by mSINGS^8^), known pathogenic *POLE* mutations^9^, and CN burden^10^. The proportion of cases in each group was similar to the previous TCGA study, except for CN-high cases reflecting the intentional enrichment of serous tumours in TCGA.

### Germline predisposition

We used WGS of constitutional DNA from these 665 cases and a further 99 cases unsuitable for tumour analysis (owing to PCR library amplification) to investigate germline pathogenic variants in Mendelian cancer genes that predispose to EC, ovarian or colorectal cancer (*MSH2*, *MLH1*, *MSH6*, *PMS2*, *POLD1*, *POLE*, *PTEN*, *APC*, *BRCA1*, *BRCA2*, *PALB2*, *TP53*, *MUTYH*, *NTHL1*, *SMAD4*, *BMPR1A, CDH1* and *STK11*). Thirty-three patients (4.3%) had previously unreported EC predisposition due to variants causal for either Lynch syndrome (LS) (19 x *MSH6*, 5 x *PMS2* including one compound heterozygote, 3 x *MLH1*, 3 x *MSH2*), Cowden syndrome (2 x *PTEN*) or Polymerase Proofreading Associated Polyposis (PPAP) (1 x *POLD1*) (**Figure 1a**). Of 24 LS carriers with tumour WGS, 23 (96%) had MSI, hypermutated tumours (range 19-432 SSV/Mb), 18 (78%) of which had detectable second hits through loss of heterozygosity (LOH) or somatic loss-of-function (LOF) mutation in MMR genes. Both Cowden syndrome cases were diagnosed with EC at a young age (30s and 50s), with tumours harbouring somatic loss-of-function (LOF) *PTEN* mutations. In addition to these cases, we identified a further 14 individuals (2.4%) with germline pathogenic variants in genes causal for cancer syndromes not known to involve EC, including *APC* (1 case), *BRCA1* (4 cases), *BRCA2* (5 cases) and *PALB2* (4 cases). The germline *APC* carrier had a somatic tumour *APC* LOF mutation, a personal history of colonic polyps, and a family history of digestive tract cancers. Of 11 cases with germline *BRCA1, PALB2,* or *BRCA2* variants and tumour WGS, 7 demonstrated somatic second hits by LOH (6 cases) or LOF mutation (1 case), with most showing high CN burden. We further investigated the association of heterozygous pathogenic *BRCA1* or *BRCA2* variants with elevated EC risk by a case/control study comparing 653 EC cases with 7,579 unrelated, cancer-free female controls of European ancestry from the 100kGP^7^. *BRCA1* variants were significantly over-represented in EC cases (RR=6.96, 95% CIs 1.67-29.08, *P*=0.02 Fisher’s exact test), though we found no evidence of an association with *BRCA2* variants (RR=1.24, 95% CIs 0.38-4.08, *P*=0.73) (**Figure 1b**). Individuals with pathogenic germline variants were significantly younger than those lacking such alterations (mean 56.6 vs 67.4 years; P=0.0071, Wilcoxon rank-sum test) (**Figure 1c**). Collectively, these data suggest that over 5% of unselected EC are caused by a germline pathogenic variant.

**Figure 1.**
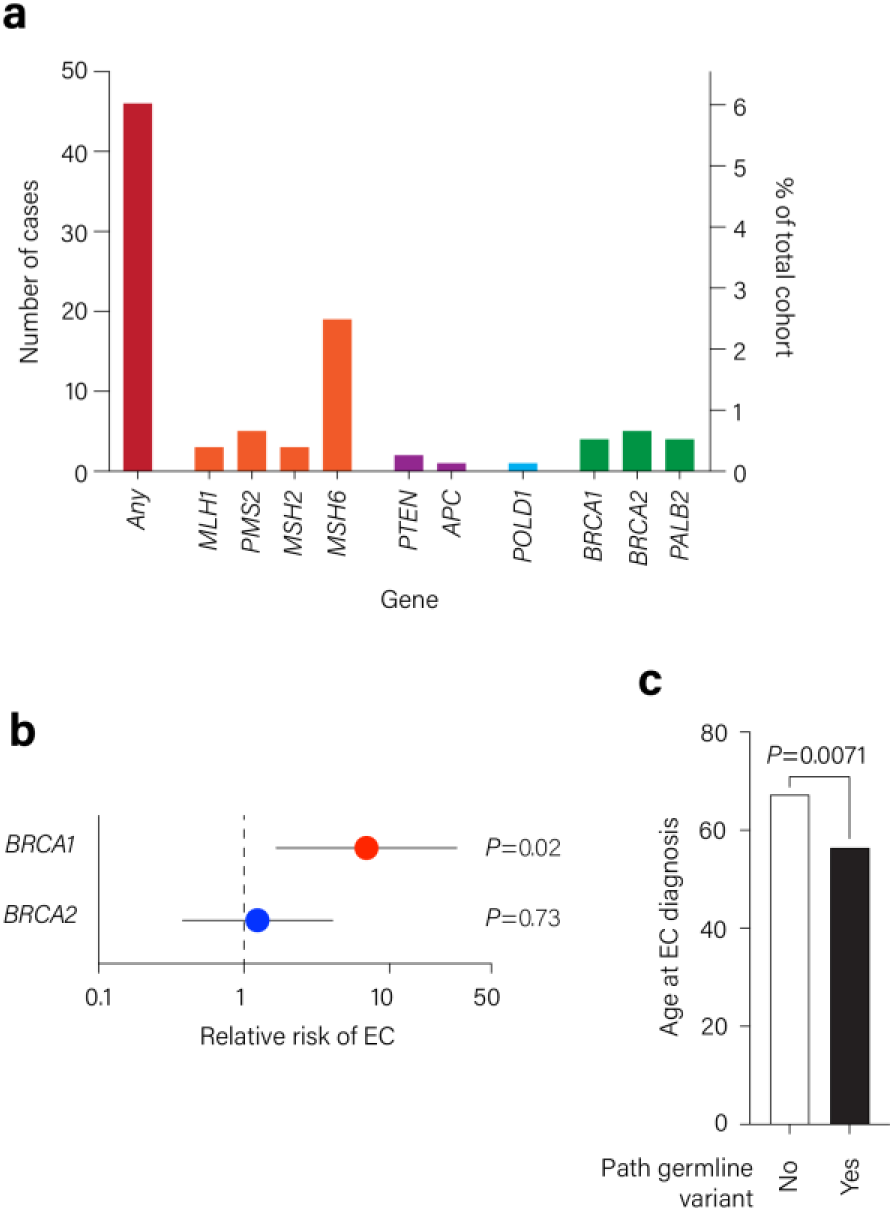
Germline cancer predisposition. **(a)** Frequency of germline pathogenic variants in genes known to predispose to endometrial, ovarian or colorectal cancer. **(b)** Relative risk of endometrial cancer associated with germline pathogenic variants in *BRCA1* and *BRCA2* determined by case-control study of 653 EC cases and 7,579 unrelated, cancer-free controls of European ancestry from the 100kGP. **(c)** Age at diagnosis of endometrial cancers in women with and without pathogenic germline predisposition variant.

### Mutational and chromosomal alteration signatures

To investigate mutational processes driving EC, we performed a *de novo* extraction of single base substitution (SBS), doublet base substitution (DBS), small (<50bp) insertion and deletion (ID), copy number (CN) and structural variant (SV) signatures^11–15^. We identified 21 SBS, 7 DBS, 8 ID, 6 CN and 9 SV signatures, most of which had been previously reported^11,14,15^. Co-occurrence of SBS10a, SBS10b, SBS28 and DBS10 with *POLE* exonuclease domain mutation (AUROC=0.905-0.998) and SBS44 with MSI (AUROC=0.94) confirmed known relationships. Mutational signatures of homologous recombination deficiency (HRD) (SBS3, ID6, ID8, and CN17), APOBEC upregulation (SBS2 and SBS13), and the tandem duplicator phenotype (SV1, CN2, CN20 and CN21), were strongly associated with CN-high subtype. Approximately one third of CN-low and CN-high tumours demonstrated high indel signature ID5, the aetiology of which is unknown. This was not associated with older age as previously reported^13^ and its biological significance is presently unclear.

### Coding drivers

We investigated coding driver genes altered by SSVs by IntOGen both across the whole cohort and within EC molecular subgroups (excluding outlier hyper-/ultramutated tumours)^16^. This identified 107 candidate EC drivers including 41 of 46 genes identified as EC drivers by ≥2 prior studies, 29 lower-confidence EC drivers (identified in a single previous study), 30 genes identified as drivers in other cancer types but not previously implicated in EC, and 7 genes with no known role in cancer (**Figure 2a**)^4,16–20^. Of the genes identified as drivers within EC molecular subgroups, six (*ARHGAP35*, *ARID1A*, *PIK3CA*, *PIK3R1*, *PTEN*, *TP53*) were identified in all subgroups, 25 in 2-3 subgroups, and 71 in a single subgroup (**Figure 2b**). Five drivers were only identified in the whole-cohort analysis.

**Figure 2.**
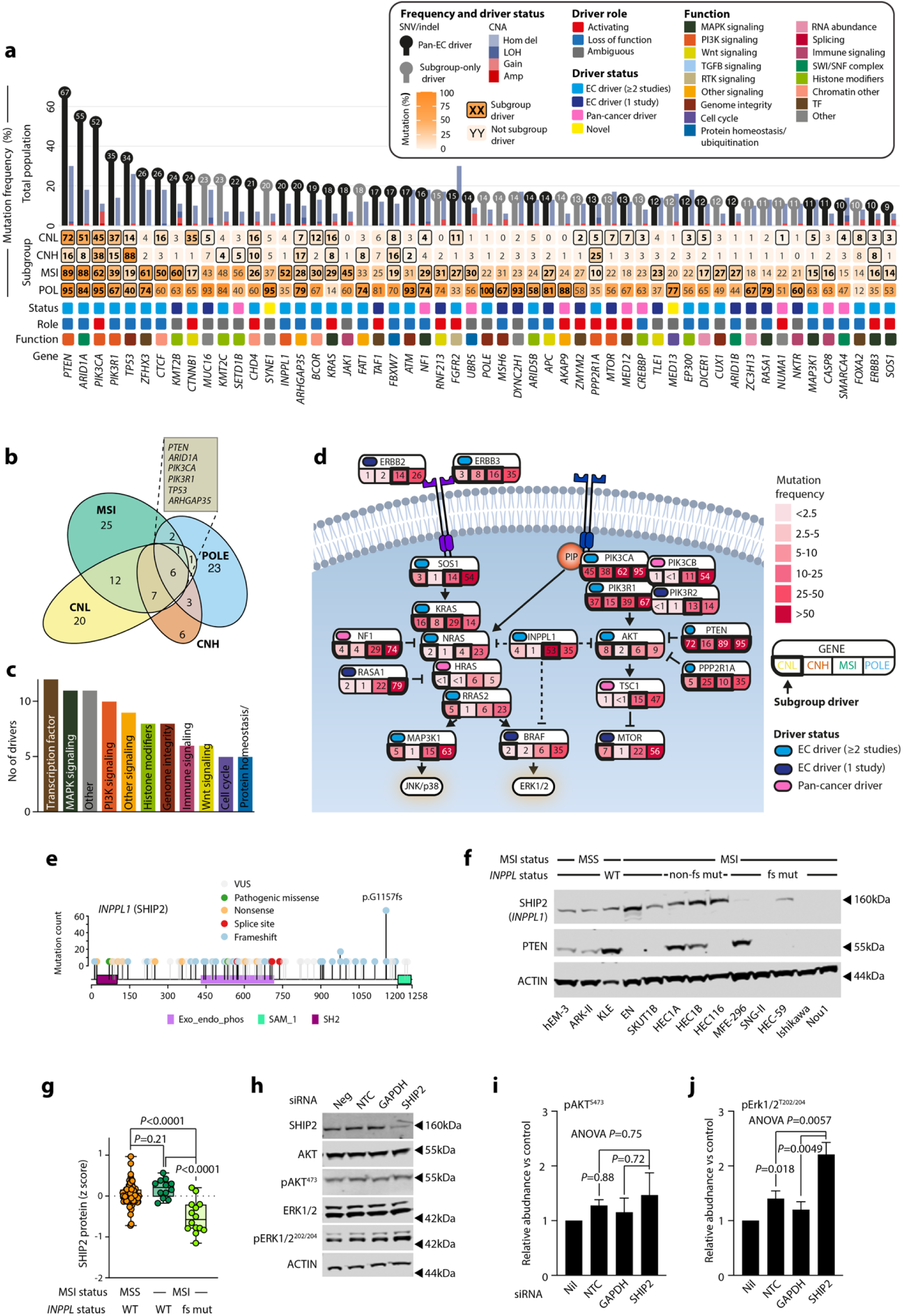
IntOGen analysis identifies known and novel endometrial cancer drivers including novel tumour suppressor *INPPL1* (SHIP2) **(a)** Top 55 most common mutational driver genes identified by IntOGen^16^ (**Methods**) in the whole cohort, and/or within EC molecular subgroups shown in descending order of mutational frequency. Functional classification is based on that used by Bailey et al^17^. Upper bar for each gene indicates oncogenic mutation frequency (both SNVs and indels), while lower bars show frequency of copy number alteration (gains, losses) at the corresponding locus. Driver role was assigned by IntOGen based on prior literature and mutation type. Prior driver status reflects whether gene was previously identified as driver/significantly mutated gene (SMG) in studies which used formal statistical methods for driver/SMG identification (i.e. excludes studies that reported recurrent mutation only in the absence of this criterion). RTK: receptor tyrosine kinase, Amp: amplification (≥5 copies), Gain: 3-5 copies, LOH: loss of heterozygosity, Hom del: homozygous deletion. **(b)** Overlap in coding drivers within EC molecular subgroups (figures indicate number of drivers in each group). The six genes identified as drivers in all four subgroups are indicated. **(c)** Number of drivers identified according to functional role after Bailey et al^17^. (**d**) Schematic showing drivers in PI3K and MAPK pathways. The frequency of oncogenic mutations within EC molecular subgroups, and the driver status from prior literature are shown. CNL: copy number low, CNH: copy number high, MSI: microsatellite instability high, POLE: *POLE* mutant. (**e**) Lollipop plots showing frequency of *INPPL1* (SHIP2) coding mutations in the whole cohort of 665 ECs. VUS: variant of unknown significance (**f**) Immunoblot showing levels of *INPPL1* protein product SHIP2 and related lipid phosphatase and prototypical EC tumour suppressor PTEN in EC cell line protein lysates according to microsatellite instability (MSI) and *INPPL1* frameshift mutation status (fs mut and non-fs mut). Actin is shown as loading control. (**g**) SHIP2 protein shown as z scores according to MSI and *INPPL1* mutational status in ECs from the CPTAC consortium^96^. Statistical significance was evaluated by unpaired ANOVA with comparison of indicated groups by Sidak’s post-test. WT: *INPPL1* wild-type. (**h**) Immunoblot showing effect of siRNA SHIP2 knockdown on downstream PI3K and MAPK pathway activation evaluated by phosphorylation of AKT at Ser 473 (pAkt^S473^) and ERK1/2 at Thr 202/204 (pErk1/2^T202/204^) respectively. Image is representative of 3-4 independent experiments (**i**,**j**) Densitometric quantification of (**i**) pAkt473 and (**j**) pErk1/2 levels as shown in (**e**) normalised to actin following siRNA against non-targeting control (NTC), positive control GAPDH, and *INPPL1*/SHIP2. Levels are shown relative to untransfected control (Nil). Error bars indicate standard error of the mean (SEM) from 3 (Akt^473^) or 4 (Erk1/2^202/204^) independent experiments. Statistical significance was evaluated by unpaired ANOVA with comparison of indicated groups by Sidak’s post-test.

Mapping drivers to biological processes (**Figure 2c**) confirmed the centrality of PI3K, MAPK and Wnt signalling dysregulation in EC, with 27 drivers mapping to these pathways, including several not previously implicated in EC such as *HRAS* and *NF1* in the MAPK pathway, *PIK3CB* and *TSC1* in the PI3K pathway, and *ZNRF3* in the Wnt pathway (**Figure 2d**). Nearly all tumours had mutations in one or more of these pathways with >25% cases carrying ≥4 alterations. Further investigation suggested intriguing relationships between drivers in the same pathway, for example significant mutual exclusivity between oncogenic mutations in PI3K pathway catalytic subunits *PIK3CA* and *PIK3CB* and regulatory subunits *PIK3R1* and *PIK3R2* (*P*=6.6e-16, permutation test). Within the PI3K pathway, we also noted frequent protein-truncating frameshift mutations in *INPPL1* (**Figure 2e**) particularly in MSI cases. *INPPL1* encodes SHIP2, a lipid phosphatase that dephosphorylates phosphatidylinositol (3,4,5)-trisphosphate (PIP3) at the 5’ position, a function analogous to *PTEN,* the PIP3 3’ phosphatase. *INPPL1* was identified as a possible EC driver by previous pan-cancer studies^17,18^, but the consequences of its mutation are unknown. Analysis of EC cell lines and ECs from the Clinical Proteomic Tumour Analysis Consortium (CPTAC) showed *INPPL1* frameshift mutations correlated with reduced or absent SHIP2 protein (**Figure 2f,g**). SHIP2 knockdown in immortalized human endometrial epithelial cells^21^ (**Figure 2h**) did not alter AKT phosphorylation (**Figure 2i**), but significantly increased activating ERK1/2 phosphorylation (**Figure 2j**), suggesting that *INPPL1* is a tumour suppressor and negative regulator of MAPK signalling.

Other cellular processes recurrently targeted by known and novel EC drivers included chromatin regulation, cell cycle control, DNA repair, RNA abundance and transcription (**Figure 2c**). Multiple drivers are known or are predicted to modify oestrogen signalling, consistent with its critical role in endometrial carcinogenesis^22^. In addition to known drivers *ESR1* and *ZFHX3*^23^, we identified previously unreported loss of function mutations in the ESR1 transcriptional repressor *SIN3A*^24^, and novel hotspot missense mutations in the ER coactivator *RBM39* ER binding domain^25^ and transcriptional regulator of oestrogen receptor expression *MED12*^26,27^. *MED12* missense mutations clustered around codon 23 rather than hotspot codons 43 and 44 mutated in uterine leiomyomas and breast fibroadenomas, suggesting distinct gain of function^26,28^. Other transcription factor drivers included *MAX*^29^, *MYCN*^30^ and *NFE2L2*, all of which showed low frequency hotspot mutations (0.8-2.0% cases).

Comparison of drivers across EC subgroups revealed notable differences. Drivers impacting oestrogen signalling were seldom mutated in CN-high cases, consistent with oestrogen independence of these tumours. *APC* was a high-frequency driver in MSI and POLE subtypes, but was seldom mutated in CN-low or CN-high tumours. Subgroup-specific drivers included *MTOR* and *UBE2A*, with uncommon hotspot mutations in CN-low tumours, *RNF213* frameshifts in MSI tumours, and *XPO1* hotspot mutations in POLE tumours.

### Non-coding drivers

Driver detection in the non-coding genome has proven challenging, with only a handful of candidates identified across cancers to date^31^. Of 97 candidate non-coding driver elements identified by OncodriveFML (Q<0.01) in our cohort, most were of unlikely pathogenicity based on their occurrence in regions of poor mappability, in highly mutated tumours, or absence of previous evidence of positive selection in cancer. Notable exceptions were non-canonical splice site mutations in *PTEN*, for which we found positive selection for 24 unique variants across 28 cases, 9 of which were predicted to have high impact by SpliceAI^32^. While *PTEN* splice site and coding mutations were not mutually exclusive, 6 of 9 high impact splice site mutations occurred in cases with *PTEN* coding mutation but no LOH, raising the possibility that they may act as second hits to coding mutations. Positive selection for non-canonical splice site and core promoter mutations in *HLX1,* a transcription factor overexpressed in leukaemia^33^, were of less certain significance as their effect on expression awaits determination. We complemented the region-agnostic analysis by manual review of known non-coding drivers^31^. *TERT* promoter mutations in 20 (3.0%) cases included known oncogenic mutations c.-124C>T (C228T) (7 cases), c.-146C>T (C250T) (1 case) and c.-57A>C (1 case)^34^, as well as c.-124C>A substitutions (3 cases), and were significantly enriched in CN-high tumours (OR = 5.16, 95% CI 1.39-20.95, Fisher’s exact test *P* = 0.0063). In contrast, variants in *TP53* 5’ region were rare (3 cases) and of uncertain pathogenicity. The scarcity of non-coding EC drivers was consistent with the earlier landmark pan-cancer non-coding driver study^31^, and supports the conclusion that these are relatively uncommon compared to coding variants.

### Copy number and structural drivers

We combined multiple methods to identify SV and CN signatures and drivers (**Methods**). SVs were called by a consensus approach with recurrent hotspots identified against a simulated empirical null distribution as previously reported^35^. After exclusion of known transposable elements, this identified 128 candidate driver SV hotspots. SV hotspots were significantly enriched at oestrogen receptor binding sites (72 of 128 hotspots, *P*=0.0082, permutation test), as previously shown in breast cancer^36^, another oestrogen-dependent malignancy, consistent with the ability of oestrogen to induce double strand breaks (DSBs)^37^. The most common recurrent SV, detected in 37 (5%) cases, was a 1.3kb inversion at chr22 which targeted an enhancer 17kb downstream of the *CHEK2* tumour suppressor, and which could plausibly impact *CHEK2* expression. Translocations within 1mb of *MYC* in 13 (2%) of tumours partnered with different regions in each case, though partner regions included known cancer genes *ESR1*, *CCNE1*, *MECOM*, *ARID5B* and several other genes with reported oncogenic potential (**Figure 3a**), suggesting possible enhancer hijacking.

**Figure 3.**
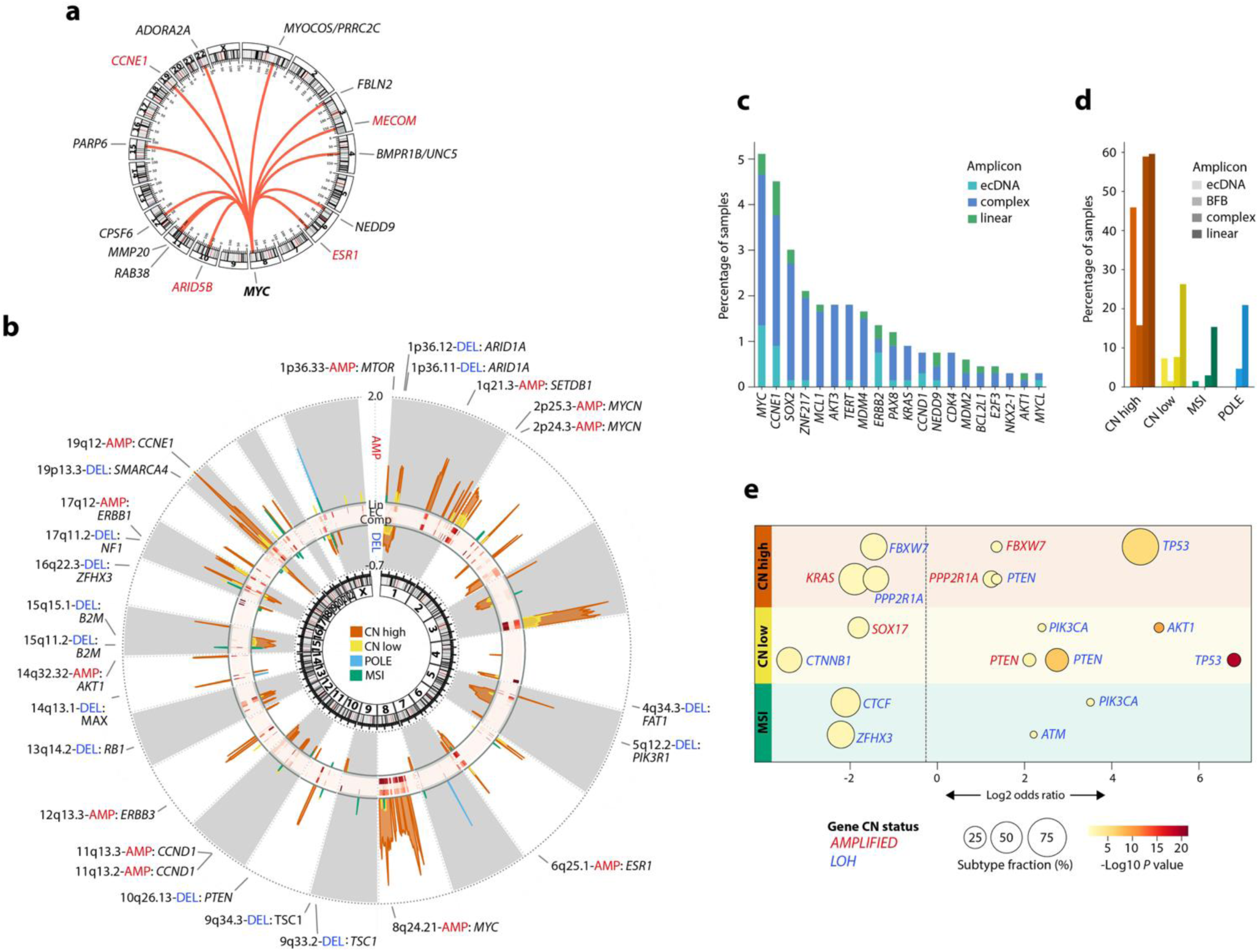
Copy number drivers and allelic imbalance. (**a**) Circos plot showing translocations involving MYC oncogene. Genes in partner regions shown with known cancer genes (included in COSMIC cancer gene census or OncoKB database) highlighted in red. (**b**) Circos plot showing GISTIC significantly recurrent focal gains and losses across EC molecular subgroups. Mechanism of amplification (classed as ≥5 copies in diploid tumours and ≥10 copies in tetraploid tumours) determined by Amplicon Architect is shown in the central bands: Lin – linear amplification, EC – extrachromosomal, Comp – complex. Annotated regions are those with GISTIC q<0.1 containing coding drivers identified by IntOgen. (**c**) Top 20 most commonly amplified genes as identified by amplicon architect. (**d**) Prevalence of amplicons due to extrachromosomal DNA (ecDNA), breakage-fusion-bridge (BFB) cycles, complex and linear amplification in the whole cohort and across molecular subgroups. (**e**) Allelic imbalance of oncogenic mutations in driver oncogenes and tumour suppressors by EC molecular subgroup. No significant associations were detected in the *POLE* mutant subgroup.

Whole genome doubling (WGD) was detected in 56 tumours, and correlated with *TP53* mutation and CN-high subgroup (*P*=3.9e-03, linear regression). Significantly recurrent chromosomal arm-level alterations in the whole cohort included frequent gains of 1q (40% cases, q=0), 21p (47% cases, q=0) and 8q (24% cases, q=0) and losses of 21p (29% cases, q=0), Xp (23% cases, q=0) and 16q (20% cases, q=0). Focal CNAs included known amplifications of gynaecologic cancer master transcription factors *MECOM*, *PAX8*, and *SOX17*^38^, p53-regulator *MDM4*, H3K9 methyltransferase *SETDB1*, and oncogenes *MYC*, *CCNE1*, *ERBB2*, *ERRB3* and *AKT1*^4,39^ (**Figure 3b**). We also confirmed focal losses of tumour suppressors *PTEN, TSC1, ZFHX3*, *CDH1* and *CTCF*^4,39^ (**Figure 3b**). Novel significantly recurrent CNAs included amplification of *MYCN* (2p25.3) – recurrently amplified in neuroblastoma^40^ – in 16% of tumours (Q_RESID_=1.7e-05), and loss of its negative regulator *NEDD4*^41^ (15q21.3) in 17% of cases (Q_RESID_=3.3e-04), with significant mutual exclusivity of the two events in CN-high cases (Log2 OR = -1.58, 95%, CI = -2.95 – -0.19, *P*=0.032). Another significantly recurrent gain of 10q11.22 in 20% of tumours contained the ER-coactivator *NCOA4*^42^ (**Figure 3b**). Further analysis by Amplicon Architect^43^ identified ecDNA as the origin of amplification of *MYC* (9 cases), *CCNE1* (5 cases), and *ERBB2* (5 cases), although these loci were more commonly amplified due to complex rearrangements (**Figure 3c**). Both ecDNA and complex amplicons were significantly enriched in CN-high ECs compared to other subgroups (**Figure 3d**).

Multiple CNAs impacted genes carrying coding drivers. Drivers showing allelic imbalance included gains of oncogenic *AKT1* and *KRAS* mutations^44^ across molecular EC subgroups, and oncogenic *PIK3CA* mutations combined with LOH in CN-low (Log2 OR= 5.08, P=9.8e-10, CI = 3.42 – 6.74) cases (**Figure 3e**). Allelic imbalance of tumour suppressors varied; while *TP53* point mutations and *NF1* LOF mutations showed loss of the wild-type allele, enrichment for gains of *PTEN* and *FBXW7* point mutations suggested either dominant negative effects or selection against gain of the wild-type allele^45,46^ (**Figure 3e**).

### Non-nuclear DNA

Alteration of the vaginal microbiome in EC has been reported^47^ but evidence for a microbial role in tumorigenesis is lacking. Bacterial genera identified in non-human reads either had no proven role in cancer (e.g. Pseudomonas), or were likely contaminants (e.g. Ralstonia and Variovorax) at low abundance compared to similar analysis of colorectal cancer^35^. Consistent with this observation, the site of sample processing was reflected in distinct clusters on UMAP of microbial species read counts, and was the strongest predictor of tumour microbiome in multivariate regression. Collectively, these results do not support a major role for the microbiome in EC tumorigenesis.

Analysis of mitochondrial DNA revealed increased mtDNA copy number in CN-high tumours, consistent with established correlations between mtDNA copy number and tumours with WGD^48^ (t-test *P*=5.53e-16, 4.85e-15 and 2.21e-13 vs CN-low, MSI and *POLE* groups respectively. In contrast to nuclear DNA, mtDNA mutation burden was significantly lower in MSI tumours compared to CN-low and CN-high subgroups (t-test *P* = 4.66e-06 and 4.52e-03 respectively). Negative selection for homoplasmic-like (variants with >80% allele frequency) truncating mutations (dN/dS = 0.18, 95% CI = 0.04-0.84, *P*=0.03) was concordant with similar analysis of other cancer types^48^. Evidence for positive selection for heteroplasmic missense mutations was inconsistent across genes and subgroups, precluding definitive conclusions.

### Immune editing and escape

We examined the immunogenomic landscape of EC. Predicted neoantigen burden varied substantially within and between EC molecular subgroups (**Figure 4a**), and positively correlated with SSV burden (*P*<10^-16^). Common EC drivers were poorly antigenic, and appeared to be selected for at the individual patient level based on their predicted binding to patient HLA molecules, consistent with prior literature^35,49^ (**Figure 4b-d**). However, *PTEN* was the only gene for which mutations were significantly more common in patients in whom they were less immunogenic (**Figure 4e**). Immune escape mechanisms differed between EC subgroups; mutations in antigen presentation genes (APG) and HLA alleles predominated in POLE and MSI tumours, while HLA LOH and allelic imbalance were predominant in CN-high tumours (**Figure 4f**). Uncommon genetic immune escape in CN-low tumours (**Figure 4g**) suggested alternative mechanisms of immune evasion. Predicted neoantigen burden was greater in MSI and POLE tumours with APG or HLA mutation and in CN-high tumours with HLA LOH, independent of SSV burden and clinicopathologic factors in multiple regression (**Figure 4g**). These results serve to highlight the role of immune editing in sculpting the EC genome and the selective pressure for immune escape in hyper- and ultramutated tumours.

**Figure 4.**
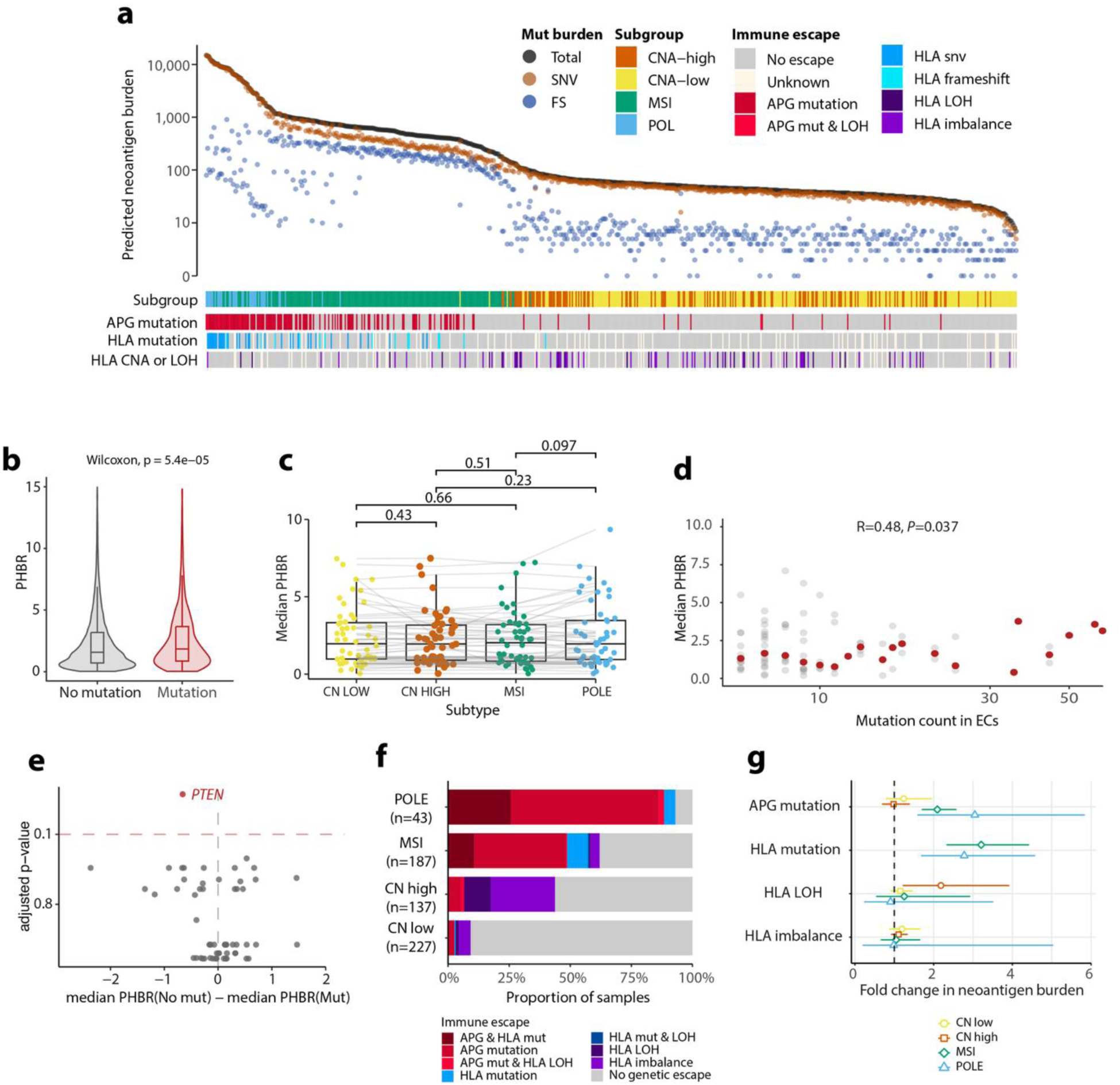
| Immunogenomic landscape of endometrial cancer. (**a**) Predicted neoantigen burden and immune escape mutations across molecular EC subgroups. Bars show antigen-presenting or antigen-processing gene (APG) and HLA alterations in each cancer. SNV: single nucleotide variant, FS: frameshift. (**b**) Predicted harmonic best rank (PHBR) of non-observed mutations in all cancers vs observed mutations. (**c**) Median PHBR of driver mutations shared across EC molecular subgroups, computed separately for tumours of each subgroup. Lines connect individual mutation PHBR values across subgroups. (**d**) Median PHBR of driver mutations across the entire EC cohort by mutation count. Grey dots represent individual mutations, red dots show the median for mutations at the same frequency. (**e**) Median PHBR difference of non-mutated and mutated driver gene alterations. Dots indicate individual drivers; those showing significant difference (*P*_Bonferroni_ < 0.1) are coloured red. (**f**) Somatic mutations in antigen presentation genes (APG) and HLA alleles by EC molecular subgroup. (**g**) Relationship between somatic immune escape alterations and predicted neoantigen burden determined by multiple regression. Symbols indicate odds ratio (OR) point estimates and whiskers 95% confidence intervals.

### Integrated analysis

We explored the relationships between mutational processes, drivers, and other alterations. Associations of coding drivers with mutational signatures mostly reflected either direct causal relationships (e.g. *POLE* mutation with SBS10a/b or *TP53* mutation with CN and SV signatures of aneuploidy) or the consequences of mutational processes (e.g. *CUX1* frameshift mutations and SBS44 secondary to MMRd). HRD signatures SBS3, ID6, ID8 and CN17 were significantly correlated with *TP53* mutation and mostly anticorrelated with mutation of CN-low drivers *PTEN* and *ARID1A*; intriguingly the anticorrelation of these drivers with SBS3 remained significant after adjusting for *TP53* mutation in multiple regression. Tandem duplicator signatures CN17 and CN20 were both strongly associated with *TP53* mutation (linear regression *P*=3.1e-10 and *P*=3.0e-25 respectively), while CN20 was also associated with *PPP2R1A* mutation (*P*=3.8e-12), CN17 was not (*P*=0.41), and neither were associated with *FBXW7* mutation (*P*=0.12 and *P*=0.61), in contrast to a previous study^50^.

Unsupervised hierarchical clustering by mutational signatures and coding drivers (**Methods**) identified known and novel tumour subgroups (**Figure 5**). The first bifurcation in the dendrogram split tumours largely according to SSV burden: an SSV high arm including three clusters, and an SSV low arm containing four clusters. Among SSV high cases, a cluster of MSI tumours with elevated indel burden and MMRd mutational signatures (SBS15, SBS26 and SBS44), and a cluster of SNV ultramutated tumours with pathogenic *POLE* mutations corresponded to TCGA MSI/MMRd and *POLE* molecular subgroups respectively. The third, smaller cluster included tumours with SBS14 (11 cases) and SBS20 (7 cases) reflective of concurrent MMRd and *POLE* or *POLD1* mutation respectively^51,52^. MSI-POLE/POLD1 tumours had SNV burden similar to the POLE subgroup, although they would be categorised as MSI/MMRd by current guidelines. Interestingly, ten (55.5%) of the MSI-POLE/POLD1 cases carried germline LS variants (i.e. 40% of 25 LS cases with tumour WGS had somatic *POLE* or *POLD1* mutations), suggesting that somatic *POLE*/*POLD1* mutations promote LS endometrial tumorigenesis, as previously shown for cancers caused by congenital MMR deficiency (cMMRd)^50^. The SSV low arm split into two branches according to tumour CN burden. An SSV low, CN low cluster included tumours with near-universal mutation of PI3K pathway genes *PTEN*, *PIK3CA* or *PIK3R1*, diploid genotype and mostly endometrioid histotype, corresponding to the CN-low subgroup in the TCGA classification (**Figure 5**). The SSV low, CN high branch included three clusters of mostly non-endometroid tumours with near universal *TP53* mutation and 17p LOH. The first of these, which we termed ‘CN-high WGD’, included cases with CN signature CN2, corresponding to whole genome duplication. The second, which we termed ‘CN-high LOH’, showed low ploidy but enrichment for signature CN9 of focal LOH (**Figure 5**). Both CN-high WGD and CN-high LOH cases showed poor outcome compared to CN-low cases (HR for overall survival = 5.4, 95% CI = 3.1-9.3, *P*<0.001; and HR = 5.0, 95% CI = 2.9-8.9, *P*<0.001 respectively). (**Figure 5**). The third, smaller CN-high cluster contained ECs with mutational signatures of HRD, six (31.5%) of which were associated with germline pathogenic variants in *BRCA1*, *BRCA2* or other HR genes (**Figure 5**). This cluster, which we termed CN-high HRD, did not show the poor prognosis associated with the other CN-high subgroups, and in fact had similar survival to CN-low cases. Thus, agnostic analysis of mutational signatures and genomic features confirmed and extended the current gold-standard molecular EC classification and revealed novel germline-somatic associations.

**Figure 5.**
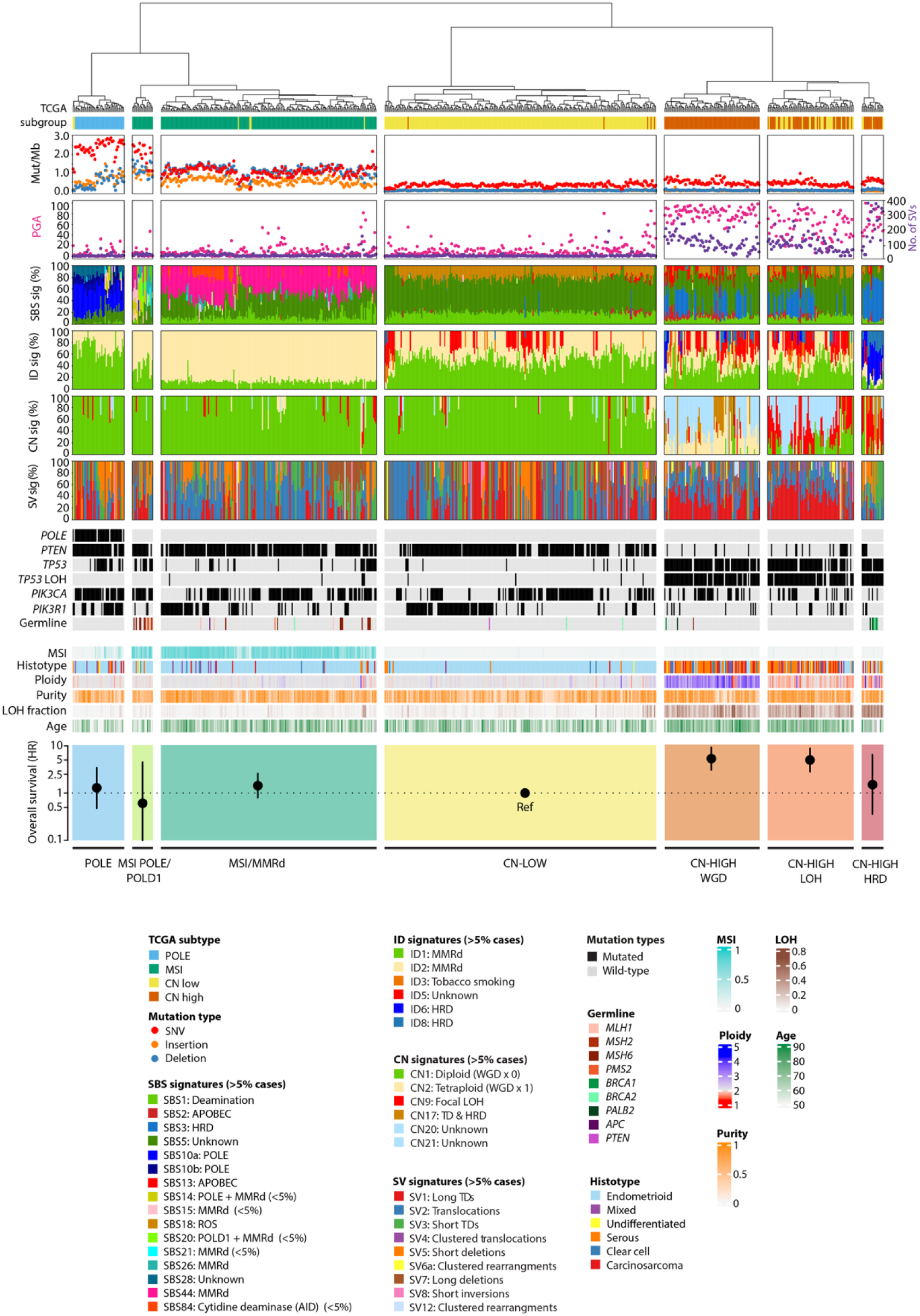
Unsupervised clustering recovers known and novel endometrial cancer molecular subtypes. Unsupervised hierarchical clustering with Laplace feature selection (see **Methods**) of tumours using single base substitution (SBS), insertion and deletion (ID), copy number (CN) and structural variant (SV) signatures and driver gene mutations as input. Mutational signatures, driver alterations, loss of heterozygosity (LOH) tumour features are shown as indicated. Hazard ratio (HR) for overall survival (OS) of clusters adjusted for age was determined by multivariable Cox regression. PGA: percentage genome altered; SV: structural variant; POLE: DNA polymerase epsilon mutated; MSI: mSINGS microsatellite instability score; MMRd: DNA mismatch repair deficiency; CN-low: copy number low; CN-high: copy number high; HRD: homologous recombination deficient.

### Prognostic value and actionability of genomic alterations

We further examined the association of tumour molecular factors with overall survival (OS) in our real-world cohort by univariable and multivariable Cox regression, adjusting for known prognostic variables including the EC molecular classification. Univariable analysis confirmed expected shorter OS with older age (uvHR=1.64 for ≥70 vs <70; 95%CI=1.37-1.97, *P*<0.001) and CN-high subgroup (uvHR=3.70 vs CN-low referent; 95%CI= 2.39-5.73, *P*<0.001) (**Figure 6a,b**). It also identified SNV and CN burden (PGA) as novel predictors of longer and shorter OS respectively, independent of age and EC subgroup in multivariable analysis (multivariable (mv)HR=0.69, 95%CI=0.50-0.95, *P*=0.025 and mvHR=1.21, 95%CI=1.06-1.38, *P*=0.005 respectively) (**Figure 6a**). Most coding drivers were not prognostic in unadjusted analysis or were prognostic owing to their association with molecular subgroups, as was the case for *PTEN* (uvHR=0.45, 95%CI=0.31-0.64, *P*<0.001 and mvHR=0.93, 95%CI=0.59-1.46, NS) and, to a lesser extent, *TP53* (uvHR=2.76, 95%CI=1.92-3.96, *P*<0.001 and mvHR=1.60, 95%CI=0.98-2.62, NS) (**Figure 6a**). However, several drivers were prognostic independent of age, subgroup, SNV burden and PGA (**Figure 6a**). Those associated with reduced survival included *FBXW7* (mvHR=1.89, 95%CI=1.18-3.02), *MYCN* negative regulator *ARHGEF12* (mvHR=2.90, 95%CI=1.18- 7.15, P=0.021), chromatin remodelers *CHD4* (mvHR=1.99, 95%CI=1.22-3.24, *P*=0.006) and *SMARCA4* (mvHR=2.21, 95%CI=1.08-4.53, *P*=0.031), and immune escape variants *B2M* (mvHR=6.46, 95%CI=1.91-21.79, *P*=0.003) and *CASP8* (mvHR=2.48, 95%CI=1.18-5.18, *P*=0.016). Drivers associated with better outcome included *PIK3CA* (mvHR=0.67, 95%CI=0.46-1.00, *P*=0.05), atypical cadherin *FAT1* (mvHR=0.44, 95%CI=0.22-0.89, *P*=0.023) and phosphatase *PPP2R1A* (mvHR=0.53, 95%CI=0.30-0.93, *P*=0.028), mutations in which have recently shown to predict immune checkpoint blockade (ICB) benefit^53^. We investigated the favourable prognosis of HRD in CN-high EC in greater detail, combining the 15 cases in the CN-high HRD subgroup with another 13 CN-high ECs with HRD mutational signatures. After confirming significantly better survival of the 28 HRD tumours compared to the 116 homologous recombination proficient (HRP) cases (Log rank *P*=0.003), we investigated the potential relationship of this to platinum chemotherapy, given the increased sensitivity of HRD tumours to such treatment. In contrast to HRP CN-high cases, in whom significantly shorter survival with platinum treatment likely reflected confounding by intention (that is, treatment selection based on high-risk features), platinum chemotherapy in HRD cases was associated with numerically improved survival, with formal testing confirming a statistically significant interaction between HRD and platinum benefit (*P*_INTERACTION_=0.045) (**Figure 6c**).

**Figure 6.**
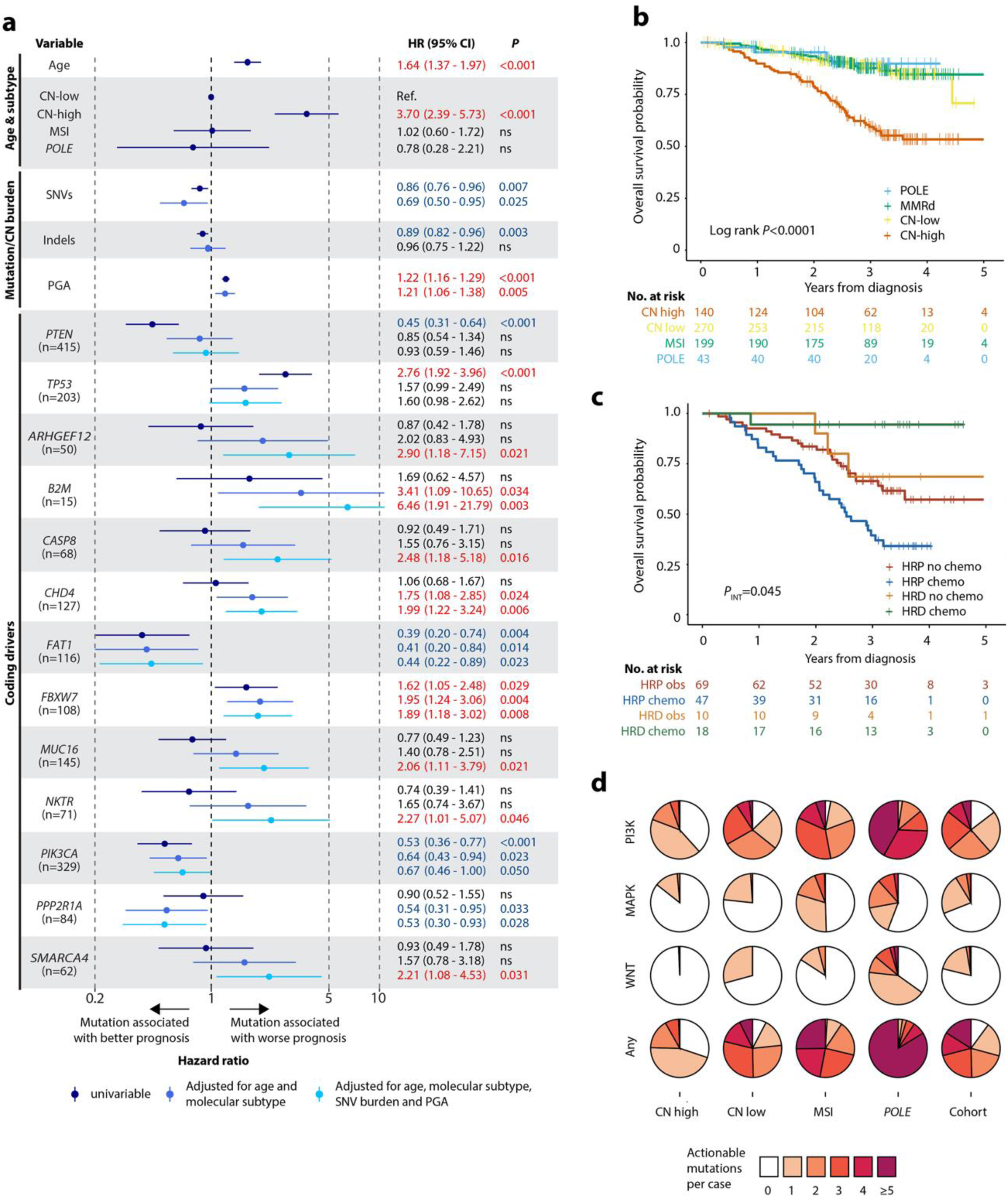
Clinical relevance of genomic alterations. (**a**) Forest plot showing univariable unadjusted and multivariable-adjusted hazard ratios (HR) for overall survival (OS) according to clinical and molecular variables. Analysis of driver mutations includes oncogenic variants only (frequency shown in parentheses). (**b**) Kaplan-Meier (KM) plot showing OS probability according to EC molecular (TCGA) subgroup. (**c**) KM plot showing OS probability of CN-high ECs according to homologous recombination deficiency (HRD) and platinum chemotherapy. (**d**) Number of actionable variants per tumour according to OncoKB and COSMIC according to signalling pathway and EC molecular subgroup. Statistical comparison between groups in **b, c** was performed by log-rank test, interaction testing in **c** was performed by Cox regression.

Referencing the OncoKB Knowledge Base (version 3.11)^54^ and COSMIC Mutation Actionability in Precision Oncology database^55^, we identified 12 drivers predictive of response to an FDA-approved or investigational drug in another cancer type (OncoKB level 1-2), and a further 10 drivers with compelling biological evidence of potential predictive utility (OncoKB level 3-4). The most common putatively actionable alterations were mutations in *PIK3CA*, *ARID1A* and *PTEN*, the frequency of which varied across molecular subgroups, as did the frequency of other actionable alterations in the PI3K, MAPK and Wnt pathways (**Figure 6d**). Other alterations detected by WGS were also recognized biomarkers of drug response. Mutational signatures of MSI/MMRd (predictive of ICB benefit), and HRD (predictive of response to PARP inhibition and platinum chemotherapy) in 202 (30.4%) and 28 (4.2%) cases respectively (with no overlap), suggested that over one-third of ECs could be candidates for these treatments. To explore the prospects of other targeted therapies being used in the same patient, we combined OncoKB clinical actionability annotations, COSMIC Mutation Actionability in Precision Oncology Product and MMR status. In total, 3,718 unique ensembl annotated targets (or 1,710 unique oncogenic targets) were present. To expand the repertoire of druggable targets, we used canSAR to map and pharmacologically annotate protein networks seeded with endometrial cancer drivers^56^. Of 308 distinct proteins identified, 201 (65%) were retrieved solely from the network analysis. Among these, 171 are targetable by existing approved therapies (including FLT3 and BCL2), while of the remaining 30, 20 have an investigational compound available and 7 possess a 3-dimensional ligandable structure. Integrating DepMap data, 22 of these proteins are predicted to be essential, and 13 have lineage-concordant specificity. Finally, we used Sylver (Synthetic Lethal Vulnerabilities Exhibiting Reciprocation)^57^, to evaluate whether tumour suppressor drivers may be candidates for synthetic lethality strategies, highlighting four relationships including with *CCND1* as a candidate knockout gene in *ATM* and *CDKN2A* deficient tumour cell lines. Collectively, these data suggested substantial opportunity for genomic-informed precision medicine in EC.

## DISCUSSION

We analysed the largest cohort of whole genome sequenced ECs to date to define the genomic landscape of this common disease. We confirmed the substantial contribution of pathogenic germline variants to EC incidence, discovered novel germline-somatic interactions, more than doubled the number of putative EC drivers identified by previous studies, confirmed tumour suppressor role of uncharacterised driver *INPPL1* (SHIP2), and identified multiple novel prognostic and targetable EC subgroups and variants. Our study extends our understanding of EC biology, and provides a valuable resource for other researchers.

Strengths of our study include its large size, real-world cohort reflective of routine practice, careful curation of clinical and pathology data by the research team at recruiting sites, deep (median 100x) WGS and comprehensive genomic analyses. Our study also has limitations. The lack of transcriptomic or single cell data, both of which were outside the scope of the 100kGP initiative, could be viewed as one. However, as DNA sequencing of EC for *POLE* mutations becomes routine practice, our results are arguably more clinically relevant than those from such methods which are currently unsuited to clinical implementation. The lack of an independent cohort for validation reflects the unique nature of this dataset; accordingly, the prognostic associations we report warrant confirmation in future studies to confirm their clinical value.

In addition to the summary results provided herein and in the accompanying raw, processed and case-level data access tables are publicly available to academic researchers via the Genomics England secure Research Environment on application (details below). We hope that many will leverage this resource to advance our understanding of EC biology, and to help improve care for patients diagnosed with it.

## AUTHOR CONTRIBUTIONS

SM: Data curation, formal analysis, investigation, methodology, validation, visualisation, writing – review & editing

BK: Data curation, formal analysis, investigation, visualisation, writing – review & editing

KK: Data curation, formal analysis, investigation, visualisation, writing – review & editing

IS: Data curation, formal analysis, investigation, writing – review & editing

EL: Data curation, formal analysis, investigation, visualisation, writing – review & editing

CAP: Data curation, formal analysis, investigation, visualisation, writing – review & editing

RC: Data curation, formal analysis, investigation, visualisation, writing – review & editing

AT: Data curation, formal analysis, investigation, writing – review & editing

LK: Investigation, writing – review & editing

YC: Investigation, writing – review & editing

AF: Data curation, formal analysis, investigation, writing – review & editing

AJC: Data curation, writing – review & editing

AH: Data curation, writing – review & editing

DC: Data curation, writing – review & editing

ASu: Investigation, writing – review & editing

BN: Data curation, formal analysis, investigation, writing – review & editing

ST: Data curation, investigation, writing – review & editing

HW: Writing – review & editing

NM: Conceptualisation, writing – review & editing

ASos: Conceptualisation, writing – review & editing

GEL: Data curation, methodology, writing – review & editing

AA: Methodology, writing – review & editing

JB: Methodology, writing – review & editing

NLB: Methodology, writing – review & editing

ASot: Funding acquisition, writing – review & editing

TB: Methodology, writing – review & editing

EC: Methodology, writing – review & editing

RE: Methodology, project administration, writing – review & editing

TG: Conceptualisation, funding acquisition, methodology, supervision, writing – review & editing

IT: Conceptualisation, funding acquisition, methodology, supervision, writing – review & editing

RSH: Conceptualisation, funding acquisition, methodology, supervision, writing – review & editing

AJG: Conceptualisation, funding acquisition, methodology, project administration, supervision, visualisation, writing – review & editing

DCW: Conceptualisation, funding acquisition, methodology, project administration, supervision, writing – review & editing

DNC: Conceptualisation, funding acquisition, methodology, project administration, resources, supervision, visualisation, writing – original draft, writing – review & editing

## ACKNOWLEDGEMENTS

E.L. is supported by Chalmers Area of Advance Health Engineering and Ragnar Söderbergs Stiftelsen.

C.A.-P. was supported by “la Caixa” Foundation (ID 100010434) fellowship (LCF/BQ/ES18/11670011).

A.Su. is supported by a Wellcome Trust Early Career Award (227000/Z/23/Z).

E.J.D. is supported by a National Institute for Health and Care Research (NIHR) Research Professorship (NIHR304303) and the NIHR Manchester Biomedical Research Centre (NIHR203308).

A.Sot. is supported by Associazione Italiana per la Ricerca contro il Cancro – AIRC (28961) and by ERC Consolidator Award (101125077).

N.L.-B. acknowledges funding from the European Research Council (consolidator grant 682398) and ERDF/Spanish Ministry of Science, Innovation and Universities – Spanish State Research Agency/DamReMap Project (RTI2018-094095-B-I00) and Asociación Española Contra el Cáncer (AECC) (GC16173697BIGA). IRB Barcelona is a recipient of a Severo Ochoa Centre of Excellence Award from the Spanish Ministry of Economy and Competitiveness (MINECO; Government of Spain) and is supported by CERCA (Generalitat de Catalunya).

T.G. acknowledges funding from Cancer Research UK (A19771), and the Wellcome Trust (202778/Z/16/Z).

I.T. is funded by Cancer Research UK grant C6199/A27327.

R.S.H. is supported by the Wellcome Trust (214388) and Cancer Research UK (C1298/A8362).

A.J.G. is supported by the University of Konstanz and a Junior Research Group (NFG027) of the Hans Böckler Foundation.

D.C.W. is supported by the NIHR Manchester Biomedical Research Centre (NIHR203308).

D.N.C is supported by a Cancer Research UK Senior Cancer Research Fellowship (RCCSCF-Nov24/100001) and previously received support from the Oxford NIHR Comprehensive Biomedical Research Centre (BRC).

We gratefully acknowledge the participants of the National Genomic Research Library (NGRL), whose contributions made this research possible. Secure access to the NGRL under project ID RR27 was provided by Genomics England, which delivers the NGRL in partnership with NHS England, and is wholly owned by the UK Department of Health and Social Care. The NGRL contains participants’ health data collected by the NHS as part of their care, along with samples and data from their participation in research, for which fully informed consent has been obtained. This includes genomic and clinical data provided through the NHS Genomic Medicine Service, as well as data obtained through research studies, including the 100,000 Genomes Project and the Generation Study, both of which are delivered in partnership with the NHS, and from other research cohorts involving external collaborators.

## CONFLICTS

BN is now an employee of Boehringer Ingelheim.

## 100kGP ENDOMETRIAL CANCER CLINICAL CONSORTIUM

### Members

Sarah Coleridge^1^, Christina Fotopoulou^2^, Amy Hawarden^3^, David Jeevan^4^, Debra Josephs^5^, Ioannis C Kotsopoulos^6^, Emma Long^7^, Neil Ryan^8^, Luke Wylie^9^, Gail Webb^1^, Lucy Wibmer^7^

### Affiliations

1. Royal Devon University Hospitals NHS Foundation Trust, UK
2. Department of Surgery and Cancer, Imperial College London, London, UK
3. Manchester University NHS Foundation Trust, UK
4. University of Birmingham, UK
5. Guys and St Thomas’ NHS Foundation Trust, UK
6. Gynaecological Oncology Department, University College London Hospital (UCLH), London, UK
7. Sheffield Teaching Hospitals NHS Foundation Trust, UK
8. University of Edinburgh, UK
9. Cambridge University Hospitals NHS Foundation Trust, UK

## Methods

### Sample collection and preparation

Clinical samples and associated data were collected as per other 100kGP studies^7,35^. Ethical approval for collection and analysis of patient samples was obtained from the HRA Committee East of England–Cambridge South Research Ethics Committee (REC reference 14/EE/1112). Individuals with suspected or confirmed endometrial cancer were identified by clinical staff at recruiting Genomic Medicine Centres. Following written informed consent, blood and tumour samples were obtained from participants, with the latter taken by specialist gynaecologic pathologists at histopathology cut-up, and frozen for later processing and downstream analysis. Adequate tumour cellularity was confirmed prior to DNA extraction by pathologist analysis of haematoxylin and eosin (H&E) stained slides. Blood and tumour samples that passed quality control underwent DNA extraction in regional genetics laboratories, with DNA transferred to the 100kGP central biorepository. WGS of paired tumour-constitutional (whole blood-derived) DNA was performed by Illumina. BAM files were transferred to Genomics England for further processing, QC and secure data storage with the Genomics England secure Research Environment. Further QC and analyses were performed by the Endometrial cancer GeCIP domain members.

### Curation of clinical data

Basic clinical and pathological data were imported from the National Cancer Registration and Advisory Service (NCRAS), run by Public Health England (PHE) and NHS Digital (NHSD) into the 100kGP secure Research Environment (RE) as part of the 100kGP project. Initial analysis of these data revealed missing or inconsistent data, particularly relating to histopathology reports. We therefore manually curated demographic and pathological data by clinicians at recruiting sites for import into the RE, substantially improving the accuracy of clinical data, and enabling us to exclude several cases wrongly classified as EC during initial data import. Survival data was obtained from the RE following import from the UK National Cancer Registry.

### Selection of EC samples analysed in this study

An initial cohort of 829 samples from the Genomics England main programme data release version 8 (v8) were exported from the Genomics England Labkey database (state January 1st 2020). From these 829 we excluded (i) formalin-fixed paraffin-embedded (FFPE) samples (68 cases) and non-FFPE samples (43 cases) which had undergone library preparation by PCR, (ii) samples with cross contamination of ≥1% of tumour or normal as estimated by VerifyBamID^58^. We next randomly selected one tumour sample for each of the six participants for whom ≥2 tumour samples were available, and then excluded cases with diagnosis other than EC, inconclusive subtype and/or missing clinical information (30 samples). We then removed five samples which were outliers across multiple metrics (percent dbSNP ids, filtered variants, ultra-low SNV burden), and a further four samples showing >25% of artefactual mutational signature SBS56. Finally, we removed the two samples from EC recurrence, leaving 665 EC samples for analysis.

### WGS and variant calling

Sequencing, alignment and variant calling used similar methods to previous 100kGP studies and were generally performed as previously described^35,59^.

### Sequencing and alignment

DNA libraries were prepared using the Illumina TruSeq DNA PCR-free kit and sequenced on a HiSeq X to generate 150 bp paired-end reads. The median sequencing depths for tumour and constitutional DNA (derived from peripheral blood mononuclear cells) were 100× and 33× respectively. Low quality outliers were identified by principal component analysis and removed based on QC metrics as follows: percentage of mapped reads; percentage of chimeric DNA fragments; average insert size; AT/CG dropout; and unevenness of local coverage. Illumina’s North Star pipeline (v.2.6.53.23) was used for the primary WGS analysis. Sequence reads were aligned to the *Homo sapiens* GRCh38Decoy assembly using Isaac (v.03.16.02.19)^60^. Outlier samples with inadequate sequencing quality were identified by principal component analysis taking into account the following quantities: (i) percent reads mapped, (ii) percent of chimeric fragments, (iii) average insert size, (iv) AT / CG dropout, and (v) local coverage divergence from a uniform distribution. Following these steps, 829 fresh-frozen, PCR-free tumour and germline WGS data from the 100kGP main programme release version 8.0 were progressed to additional filtering (**Figure S1**) and downstream analysis.

### Germline pathogenic variants

Whole genome-sequenced constitutional DNA was analysed for germline mutations in Mendelian cancer predisposition genes (*MSH2*, *MLH1*, *MSH6*, *PMS2*, *POLD1*, *POLE*, *PTEN*, *APC*, *BRCA1*, *BRCA2*, *PALB2*, *TP53*, *MUTYH*, *NTHL1*, *SMAD4*, *BMPR1* and *STK11*). Protein-truncating variants (PTVs) and missense mutations classified as ‘pathogenic’ or ‘likely pathogenic’ by ClinVar^61^ were considered as disease-causing changes, with exception of *POLE* and *POLD1* variants, which were identified as described below. Evidence of pathogenic biallelic *MUTYH* or *NTHL1* mutations was required to diagnose the recessive conditions caused by these variants, but was not found in any case. Two cases with possible germline *TP53* variants were found to have lower-than-expected variant allele frequencies (VAF), with further investigation demonstrating one likely due to postzygotic mosaicism (PZM) and the other to clonal haematopoiesis of indeterminate potential (CHIP). Supporting evidence for a causal role of germline pathogenic variants in tumorigenesis was investigated by interrogation of associated tumours for second hits, including somatic pathogenic variants or loss of heterozygosity at the affected locus.

### Somatic single-nucleotide variant and indel calling

Insertions and deletions (Indels) and single nucleotide variants (SNVs) were called using Strelka (v2.4.7)^62^. In addition to the default Strelka filters, we excluded variants based on the following parameters:

- **Poor mappability variants:** variants located in regions where the majority of overlapping reads are not uniquely mapping to the variant position.
- **Variants with a frequency smaller 5% in the 100kGP cancer dataset:** Based on the frequency of recurrent non-synonymous variants in Cancer Gene Census genes^55^ the cut-off of 5% was chosen.
- **High sequencing noise insertions and deletions:** indels in regions where greater 10% of the base calls in a window extending 50 base pairs on either side of the insertion/deletion were filtered out by Strelka due to poor quality.
- **GnomAD variants:** variants with a population germline allele frequency greater 1% in gnomAD^63^.
- **Insertions and deletions in the vicinity of Gnomad variants:** Indels that are located within 10 base pairs of 100kGP or Gnomad germline indels with allele frequency greater 1%.
- **Potential germline variants:** variants with a germline allele frequency greater 1% in the full 100kGP dataset.
- **Mapping and calling artefact variants:** SNVs that potentially result from systematic mapping and calling artefacts present in both normal and tumour 100kGP sample sets. The ratio of tumour allele depths at each somatic SNV site was tested for being significant different to the ratio of allele depths at this site in a panel of normal samples (PoN) using Fishers exact test. The PoN was created based on a cohort of 7,000 non-tumour genomes from the 100kGP dataset. At each genomic site only individuals not carrying the relevant alternative allele were included in the count of allele depths. The bcftools (v1.9) mpileup function was used to count allele depths in the PoN. To replicate Strelka filters duplicate reads were removed and quality thresholds set at a mapping quality greater or equal 5 and base quality greater or equal 5. All somatic SNVs with Fishers exact test phred score smaller 80 were filtered, with the threshold determined by optimising precision and recall calculated from a TRACERx truth set^64^.
- **Tandem repeats variants:** variants overlapping with simple repeats as defined by Tandem Repeats Finder^65^.

### Removing alignment bias introduced by soft clipping of semi-aligned reads

The *–clip-semialigned* parameter of the Isaac aligner^60^ performs the soft clipping of sequencing read ends until five consecutive bases are matching the used reference genome. Thus, the soft clipping process results in the loss of support for alternate alleles positioned within five bases of sequencing read ends. The consequence are artefactually low Variant allele frequencies (VAFs). To remove these introduced allelic bias *FixVAF* was used to soft clip all sequencing reads by five bases at the end^66^ as performed previously^35^.

### Mutational signatures

Single-base substitution (SBS), doublet-base substitution (DBS), as well as insertion and deletion (ID) signatures were extracted *de novo* and decomposed to known mutational signatures (COSMIC version 3.2) using SigProfilerExtractor^67^.

### Calling structural variants

Structural variants were identified using a similar approach to that previously reported^35^. Somatic rearrangements were called using a graph-based consensus approach including Delly^68^, Lumpy^69^ and Manta^70^ with support from CNAs. Rearrangements were initially called using the three individual callers using default parameters. Rearrangements thus identified were then filtered to exclude those (i) in which reads supporting the variant were also identified in the matched normal; (ii) where <2% of tumour reads supported the variant, or (iii) if either variant breakpoint was in a telomeric or centromeric region or on a non-standard reference contig (that is, not chromosomes 1–22, X or Y). Remaining post-filter rearrangements were merged with a modified version of PCAWG Merge SV, which uses a graph-based approach to identify and merge rearrangements identified by different callers, which permits a 400 bp difference in breakpoint position to account for variant calling ambiguity^71^. Rearrangements were passed for inclusion either if they were identified by ≥2 callers, or by a single caller with a breakpoint <3 kb from a CNA segment boundary. SVs were called in 637 out of 665 samples with CNA profiles passing quality control criteria.

We characterised genomic regions enriched for structural variations (SVs)—including deletions, duplications, inversions and translocations—using SV calls obtained from our samples. To evaluate associations between genomic context and SV frequency, we fitted negative binomial regression models incorporating SV class, mutation size, the number of mutations observed at each genomic coordinate, and the number of samples in which each mutation was detected. SVs were subsequently simulated to generate null expectations for SV burdens across the genome, preserving both the number and length of observed events. The genome was partitioned into non-overlapping 1-Mb bins, and genomic features were normalised to estimate the expected break-end frequency per bin. For each SV class, 1,000 simulated datasets were generated to derive empirical null distributions, excluding centromeric and telomeric regions. SV hotspots were identified using piece-wise constant fitting (PCF). Inter-mutational distances were log-transformed, segmented, and compared with expected break-end densities derived from the simulated datasets. False discovery rates (FDRs) were estimated from the distributions of observed and simulated segments. PCF parameters for hotspot detection were set at γ = 10 and kmin = 4 in conjunction with FDR thresholds. Hotspots lacking supporting copy-number alterations were considered artefactual and were excluded.

A permutation test was used to evaluate whether the observed overlap of SV hotspots with ER binding sites exceeded that expected by chance. To generate a null distribution, 130 random genomic locations (excluding chromosome Y, telomeres and centromeres) were selected and assigned SV lengths by randomly shuffling the lengths of the observed SV hotspots. For each iteration, the number of simulated SV hotspots overlapping ER binding sites was recorded. This procedure was repeated 10,000 times, and a *P* value was calculated as the proportion of permutations producing an equal or greater number of overlaps than observed.

Retrotranspositions were not considered for analysis in this study.

### Calling copy number alterations

Somatic CNA calling was done as previously reported^35^, using R package CleanCNA. Genome-wide subclonal CNAs were first called using Battenberg (v.2.2.8)^72^. CNA calling quality was checked using DPClust^72^ and CNAqc^73^ to CNA profiles and SNV VAFs. Samples were classified as ‘pass’ if they met both of the following criteria:

1. A clonal cluster of SNVs (0.95 ≤ CCF ≤ 1.05) as identified by DPClust. This clonal cluster was required to have either the highest CCF of all SNV clusters or contain the largest number of SNVs. SNV clusters containing <1% of all sample SNVs were removed before assessment.
2. A difference in purity estimates from Battenberg and CNAqc of <5%. CNAqc estimates sample purity considering peaks in SNV VAF distributions in genome regions with one of five copy number states (1:0, 1:1, 2:0, 2:1, 2:2).

Samples not meeting both these criteria were excluded. Individual samples underwent CNA profiling up to four times per sample, with the procedure stopped if both criteria were met. After a failure, CNA were re-called using Battenberg with re-estimated sample purity and tumour ploidy. After the first fail, purity and ploidy were re-estimated using DPClust as previously^35^. Samples failing a second time had purity and ploidy re-estimated by Ccube^74^, and those failing a third or fourth time underwent purity and ploidy estimation by CNAqc. Samples that failed after four re-runs were removed from downstream analyses dependent on CNAs. Pass CNA profiles were produced for 498 out of 665 samples. The percentage of genome altered (PGA) was defined as the percentage of genome deviating from copy number status 1/1 in tumours in which the fraction of the genome with 1/1 status was greater than that with 2/2 status, and as the percentage of genome deviating from 2/2 otherwise.

### Detection of recurrent broad and focal copy number alterations

#### Preparing input copy number segmentation file

For every tumour passing copy number quality control measures, a copy number segmentation file was generated as input for GISTIC from Battenberg per-tumour segmentation output. From each copy number segment identified by Battenberg, the chromosomal coordinates, major (nMaj) and minor (nMin) copy number calls were obtained. In the case of subclonal copy number segments, nMaj and nMin were taken from the subclone with the largest tumour fraction.

Per-segment normalised copy number was calculated differently depending on whether the tumour had been identified as having undergone whole genome doubling (WGD) where the ploidy was assumed to be 4 (tetraploid), otherwise the ploidy was assumed to be 2 (diploid).

For diploid tumours per-segment normalised copy number was calculated by:

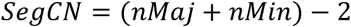

For tetraploid tumours per-segment normalised copy number was calculated by:

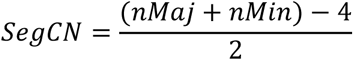

*SegCN* was thresholded to a maximum of 2.0 and minimum of -2.0.

#### Running GISTIC

GISTIC v2.0.2.3^75^ was run to identify recurrent arm-level copy number events, as well as focal amplifications and deletions (https://github.com/broadinstitute/gistic2). The following parameters were used:

“-conf 0.99 -broad 1 -qvt 0.25 -genegistic 1 -gcm extreme -brlen 0.5 -rx 0 -twoside 1 -scent median -armpeel 1 -arb 1 -refgene hg38.UCSC.add_miR.160920.refgene.mat“

### Prioritising likely gene targets of focal amplifications and deletions

Candidate target genes at focal amplifications and deletions were annotated according to the following criteria:

1. Overlap with genes at focal amplifications and deletions previously reported in a pan-cancer TCGA study as found in Supplementary Table 3 of Zack, et al. ^39^ Comparisons were made both with the overall pan-cancer GISTIC analysis, as well as GISTIC analysis restricted to the given tumour type. Special consideration was given to genes specifically highlighted in the Zack et al., 2013 analysis as being prioritised candidates
2. Overlap with Cosmic Cancer Gene Census genes and whether their annotated role (Oncogene (OG)/tumour suppressor gene (TSG)/ambiguous) is consistent with the copy number change (OG with amplifications, TSG with deletions)
3. Overlap with driver genes identified as being significantly mutated in this study, and whether the driver’s likely role (Oncogene (OG)/tumour suppressor gene (TSG)/ambiguous) is consistent with the copy number change (OG with amplifications, TSG with deletions)

On the basis of the above criteria, “consensus” driver genes were manually assigned to peaks. To maximise detection of overlapping genes, comparisons were made with all potential gene synonyms as available from the HUGO gene nomenclature name committee (https://www.genenames.org/).

### Extrachromosomal DNA detection

Potential extrachromosomal DNA (ecDNA) molecules were detected from tumour bam files using AmpliconArchitect v1.2^43^ (https://github.com/virajbdeshpande/AmpliconArchitect). Briefly, per-tumour “seed” regions were prepared from Battenberg copy-number segmentation output if a segment was >100kb and the total copy number was > 5. AmpliconArchitect was then run using these “seed” regions to extract overlapping sequence reads from the tumour bamfile and construct candidate amplicons.

Candidate amplicons were classified using AmpliconClassifier v0.4.6 (https://github.com/jluebeck/AmpliconClassifier) into the following categories: 1) Cyclic (truly circularised ecDNA) 2) Complex non-cyclic 3) Linear amplification 4) No amp/invalid. Amplicons were highlighted if containing a known highly amplified oncogene (*MDM2*, *MYC*, *EGFR*, *CDK4, ERBB2, SOX2, TERT, CCND1, E2F3, CCNE1, CDK6, MDM4, NEDD9, MCL1, AKT3, BCL2L1, ZNF217, KRAS, PDGFRA, AKT1, MYCL, NKX2-1, IGF1R, PAX8*; as per Kim et al^76^).

Genome-wide copy-number alterations were visualised using the python library pyCirclize (v.1.10.1) based on the GISTIC output (Figure 3b). For each subtype, significantly amplified and deleted regions were extracted and corresponding GISTIC g-scores were plotted. Additionally, amplified regions detected by AmpliconArchitect as either linear, complex or ecDNA were integrated by computing relative prevalences and scaling values 0 to 1.

### Classification of molecular subtypes

Somatic variants in *POLE* were classed as pathogenic based on prior literature^9^. Confirmation that all cases with pathogenic *POLE* mutation demonstrated tumour ultramutation and *POLE* SBS signatures SBS10a/b confirmed the validity of this approach. Tumour microsatellite instability was identified using mSINGS^8^ as previously described^35^. MSI/MMR status was not reliably available for ECs included in this cohort, as recruitment predated guidance recommending MMR testing in routine practice^77^. We therefore used a set of 132 colorectal cancers of known MSI status to train a model which perfectly distinguished MSI and MSS samples in a test set of CRCs using default MSINGs settings^35^. Classification of ECs as MSI by this model showed near-perfect correspondence with the presence of known MSI-associated single base substitution (SBS) signatures 44, 14 and 20, confirming its accuracy. Classification of POLE-wild-type, MSS cases into copy number high and copy number low subgroups was performed by dichotomisation of cases at the minimum of the bimodal distribution obtained from a kernel density plot of the percentage genome aberrant (PGA) of tumours – that is, MSS cancers with PGA greater than the cutpoint were considered CNA-high, whereas all other MSS cancers were considered CNA-low. Intersection of molecular EC subgroups with histopathological subtype, *PTEN* and *TP53* mutation status confirmed expected enrichments across subgroups.

### Single nucleotide variant and indel drivers

Single nucleotide variant (SNV) and insertion-deletion (indel) mutational drivers were identified using the IntOGen pipeline as reported previously^35^. Somatic mutations were annotated to Ensembl (v.101, GRCh38) using Variant Effect Predictor (VEP) using the following parameters used: vep -i <INPUT_VCF> --assembly GRCh38 –no_stats –cache –offline –symbol –protein -o

<OUTPUT> --vcf –canonical –dir <REF_DIR> --hgvs –hgvsg –fasta <GRCH38_FASTA> --plugin CADD,<CADD_SCORE_FILE> --plugin UTRannotator,<GRCH38_UORF_REFERENCE>. The CADD score file was obtained using CADD (v.1.6)^78^, and scores obtained for all SNV and indel mutations using the CADD software available from GitHub (https://github.com/kircherlab/CADD-scripts) before being utilized by the VEP CADD plugin.

#### Protein-coding driver identification

Protein-coding driver genes were identified in the whole cohort, and EC molecular subgroups individually by IntOGen (v.2020, downloaded February 2021)^16^. Background mutation rates and spectra were corrected for in each case, and mutations present in a Hartwig Consortium panel of controls^79^ were excluded from analysis. Outlier hypermutated tumours with >10,000 mutations and mutation count of greater than the upper quartile + (1.5 × interquartile range) were excluded from driver gene identification. Unless otherwise specified, mutations were mapped to canonical protein-coding transcripts from Ensembl (v.101, GRCh38).

#### Driver identification methods

The IntOGen pipeline includes seven methods of driver gene identification:

1. dNdSCV (v.0.1.0)^18^ detects mutated genes under positive selection based on an excess of non-synonymous (missense, nonsense, essential splice) mutations after correction for local trinucleotide context. The parameter ‘max_coding_muts_per_sample = Inf’ was used in the *POLE* subgroup owing to the high proportion of hypermutated tumours.
2. OncodriveFML (v.2.4.0)^80^ detects driver genes based on an enrichment of mutations of high functional impact (CADD scores^78^).
3. OncodriveCLUSTL (v.1.1.3)^81^ detects driver genes enriched for linear mutation clusters. In the primary POLE cohort, pentamer signatures were preferred to trinucleotide signatures owing to superior performance of pentanucleotide-based background models in this subgroup.
4. cBaSE (v.1.1.3)^82^ detects driver genes under positive selection based on a significant mutation count bias following correction for trinucleotide context.
5. MutPanning (v.2)^83^ detects driver genes based on enrichment of mutations with unusual nucleotide contexts compared with a background model.
6. HotMaps3D (v.1.1.3)^84^ detects driver genes based on identification of missense mutations spatially co-clustered in the protein three-dimensional structure. Protein structures were downloaded from The Protein Data Bank^85^ (downloaded March 2020).
7. smRegions (v.1^86^) detects genes containing an enrichment of non-synonymous mutations in regions of interest, such as protein domains, after correcting for trinucleotide context. This analysis utilized information from protein family (Pfam) domains that were mapped to Ensembl (v.101) canonical transcripts.

#### Combination of driver identification methods

Output from the seven driver identification methods were combined as previously reported^35^. The driver combination procedure considered the top 100 ranked genes and their associated *P* and *Q* values in each method. Somatically mutated genes assigned as tier 1 or tier 2 in the COSMIC Cancer Gene Census (CGC; v.92)^20^ were designated as the truth set of known drivers. Through comparison of the relative enrichment of CGC genes in the top ranked gene lists, a per-method weighting was obtained. Per-method ranked lists were combined using Schulze’s voting method to generate a consensus ranking, with combined *P* values estimated using a weighted Stouffer *Z* score method. Driver candidates were then classified into the following tiers:

- Tier 1: candidates for which the consensus ranking was higher than the ranking of the first gene with Stouffer *Q* ≤ 0.05. These represent high-confidence drivers.
- Tier 2: candidates not meeting the criteria for tier 1, but which are CGC genes and showed a combined Stouffer *Q*_CGC_ < 0.25. These represent a set of ‘rescued’ known cancer drivers.
- Tier 3: candidates not meeting the criteria for tier 1 or tier 2 but with Stouffer *Q* < 0.05. These represent lower confidence drivers.
- Tier 4: candidates not meeting criteria for tier 1 or tier 2 and with Stouffer *Q* > 0.05. These genes are not likely to be drivers.

### Post-processing of candidate drivers

Candidate driver genes were filtered based on the following annotations:

- Automatic fail: a candidate driver gene would be excluded from further consideration if annotated with at least one of the following:

- Tier 4: categorized as tier 4 by the combination procedure.
- Single method: only significant (*Q* < 0.1) in one of the seven methods (non-CGC genes).
- Expression: gene has very low or no expression in endometrial cancer based on data from The Cancer Genome Atlas (TCGA).
- Olfactory receptor: gene is in list of olfactory receptor genes.
- Known artefact: gene is in a list of known artefacts or long genes (for example, TTN).
- Manual review: if a gene is not excluded based on any automatic fail filters, it is retained as a candidate driver:

- Germline: non-tier 1-CGC gene has ≥1 mutations per sample and oe_syn/ms/lof > 1.5 based on gnomAD (v.2.1) constraint metric estimates.
- Sample 3 Muts: non-CGC gene for which there are ≥3 mutations in ≥1 tumour.
- Literature: non-CGC gene for which there are no literature annotations according to CancerMine
- Automatic pass: is not flagged by any automatic fail or manual review filters.

Candidate driver roles were assigned on the basis of dN/dS ratios for missense (wmis) and nonsense (wnon) mutations for the given gene derived from dNdSCV (https://bitbucket.org/intogen/intogen-plus/src/master/core/intogen_core/postprocess/drivers/role.py):

- A distance metric was calculated by distance = ((wmis – wnon))/√2
- Candidate drivers with distance >0.1 represent those with an excess of missense to nonsense mutations and are therefore considered oncogenes.
- Candidate drivers with distance <0.1 represent those with an excess of nonsense to missense mutations and are therefore considered TSGs.
- Otherwise, the role of the candidate driver is unclear and considered ambiguous.

In the case of multiple cohorts being run representing subsets of a given tumour type, a consensus role was designated comparing between each subtype role:

- Oncogene if assigned as oncogene in ≥1 cohort and as TSG in no other cohort.
- TSG if assigned as TSG in ≥1 cohort and as oncogene in no other cohort.
- Ambiguous otherwise.

Gene candidates were classified as known EC drivers, known drivers in cancer types other than EC, or potential novel drivers according to their previous identification in studies which used formal statistical methods for driver identification^4,16–20^. Prior reports of recurrent mutation alone was regarded as insufficient evidence of driver status.

#### Non-coding driver identification

##### Defining sets of non-coding regions

The following sets of non-coding regions were defined. Note that regions from non-canonical splice sites, 5’ and 3’ untranslated regions (UTRs), core and distal promoters that overlapped coding sequences (CDS) were removed using bedops (v.2.4.39)^87^. Similarly, exonic regions from canonical protein-coding transcripts were removed from long intergenic non-coding RNAs (lincRNAs), microRNAs (miRNAs), enhancers, open chromatin regions, CTCF sites and transcription factor binding sites:

- Non-canonical splice regions (*n* = 18,163). Regions extending 30 bp into the intron from essential splice donor or acceptor sites in canonical protein-coding transcripts.
- 5′ UTRs (*n* = 18,613). Regions defined based on canonical protein-coding transcripts.
- 3′ UTRs (*n* = 18,806). Regions defined based on canonical protein-coding transcripts.
- Core promoters (*n* = 19,283). Regions spanning 200 bp 5’ (upstream) and 50 bp 3’ (downstream) of transcription start sites (TSS) of canonical protein-coding transcripts.
- Distal promoters (*n* = 19,296). Regions spanning 2 kb 5’ (upstream) the TSS of canonical protein-coding transcripts.
- lincRNAs (*n* = 16,510). Exonic regions from transcripts annotated as lincRNAs in Ensembl (v.101).
- miRNAs (*n* = 1,793). Regions from transcripts annotated as miRNAs in Ensembl (v.101).
- Enhancers (*n* = 130,996). Regulatory elements annotated as ‘enhancer’ in Ensembl (v.101).
- Open chromatin regions (*n* = 95,344). Regulatory elements annotated as ‘open chromatin’ in Ensembl (v.101).
- CTCF sites (*n* = 173,711). Regulatory elements annotated as ‘CTCF sites’ in Ensembl (v.101).
- Transcription factor-binding sites (*n* = 29,259). Regulatory elements annotated as ‘TF binding sites’ in Ensembl (v.101).

#### Detecting non-coding drivers

Potential non-coding driver mutations were investigated across EC molecular subgroups. OncodriveFML (v.2.4.0)^80^ was run on sets of non-coding regions to detect biases in the functional impact of somatic mutations. OncodriveFML was run scoring indel functional impact as the maximum effect among all possible substitutions and simulating indel background models as substitutions. Elements with *Q* < 0.01 were considered as significant.

#### Analysis of splice site mutations

Mutations overlapping *PTEN* splice sites were assessed with SpliceAI^32^ and Pangolin^88^ at spliceailookup.broadinstitute.org Mutations were assigned to a splicing alteration type (Acceptor Loss, Donor Loss, Aceptor Gain or Donor Gain) based on the top delta score alteration type from SpliceAI and Pangolin followed by manual review. Mutations with both SpliceAI delta score > 0.5 and Pangolin delta score > 0.5 were classified as high impact.

#### SNV mutations exhibiting extreme strand bias

SNV mutations that otherwise passed filtering criteria as previously detailed were further scrutinized for excessive strand bias (Strelka INFO field SNVSB > 10). This highlighted many mutations that cause a recurrent missense change in *CACNA1E* (p.Ile95Leu); these exhibited excessive strand bias and were therefore deemed false calls.

### Driver mutation annotation

Non-synonymous mutations in the 682 gene transcripts considered by OncoKB (v.3.3) were annotated using the OncoKB API^54^. HGVSg identifiers were initially used; in the rare instances that this failed, a combination of gene symbol, consequence and HGVSp were used to map mutations.

#### OncoKB annotation

Non-synonymous mutations in candidate driver genes were annotated as pathogenic if any of the following criteria were met:

- The mutation is annotated by OncoKB as ‘oncogenic, ‘likely oncogenic’ or ‘predicted oncogenic’.
- The driver is classified as an oncogene, the mutation consequence is missense, and the mutation is recurrent (seen in ≥3 tumours in cohort).
- The driver is classified as a TSG or ambiguous and either:

- Consequence is protein-truncating (splice acceptor, splice donor, frameshift, stop lost, stop gained or start lost).
- Consequence is missense and mutation is recurrent (seen in ≥3 tumours in cohort).

For *POLE*, oncogenic annotations were restricted to missense mutations in the exonuclease domain (amino acid residues 268–471). Non-synonymous mutations not meeting these criteria were considered as variants of uncertain significance.

### Lollipop plots of driver gene mutations

Lollipop plots of driver gene mutations were generated using the Rpackage trackViewer^81^. Pfam protein domains mapping to the Ensembl (v.101) canonical transcripts were plotted. The protein position was taken from the first position in the HGVSp annotation, apart from splice donor and acceptor mutations, for which the codon nearest to the HGVSc transcript position was assigned as the protein position.

### Immune analysis

HLA typing of blood-derived normal samples was performed using HLATyper from the Illumina Whole Genome Sequencing Service informatics pipeline. For each sample, the top-ranked, best-supported allele pair for each type-I HLA gene locus (HLA-A, HLA-B, and HLA-C) was selected, resulting in a six-allele HLA genotype per sample. To determine the immune escape status, we evaluated the three following distinct mechanisms of (i.) somatic mutation of HLA genes, (ii.) loss of heterozygosity (LOH) at the HLA locus, (iii.) mutation or LOH affecting other antigen-presentation genes (APGs). Somatic mutations in HLA genes were identified using POLYSOLVER. Previously determined HLA genotypes were first converted into a format compatible with POLYSOLVER (lowercase, digits separated by underscores) and written to patient-specific winners.hla.txt files. The POLYSOLVER mutation-calling pipeline (shell_call_hla_mutations_from_type) was then applied to matched tumour-normal samples to detect tumour-specific alterations in HLA-aligned reads using the MuTect variant caller. Additionally, Strelka was used to identify short insertions and deletions, providing increased sensitivity compared with POLYSOLVER’s default variant caller. All SNVs and InDel variants passing quality control were subsequently annotated using POLYSOLVER’s annotation script (shell_annotate_hla_mutations). LOH status at the HLA loci was inferred using LOHHLA. Again, the winners.hla.txt HLA typing files were used as input, together with POLYSOLVER’s comprehensive, deduplicated FASTA reference of HLA haplotypes. Type I HLA alleles were classified as exhibiting allelic imbalance (AI) if the *P* value for unequal allele support was <0.01. Alleles showing AI were further classified as LOH when all the following criteria were met:

1. Estimated copy number of the lost allele was <0.50 with an upper confidence interval <0.70
2. Copy number of the retained allele exceeded 0.75
3. More than 10 mismatched sites were present between the two alleles

Overall, 98 samples had one or more haplotypes incompatible with POLYSOLVER, resulting in HLA mutation and LOH calling performed only on the compatible haplotypes (1, 2, and 3 haplotypes were excluded in, 73, 22, and 3 samples, respectively). Additionally, LOH was not assessed for 2 samples who were homozygous at all type-I HLA genes. We also considered somatic mutations and copy number alterations in the following APGs: *B2M*, *CALR*, *CANX*, *CIITA*, *ERAP1*, *ERAP2*, *HSPBP1*, *IRF1*, *PDIA3*, *PSMA7*, *PSME1*, *PSME2*, *PSME3*, *TAP1*, and *TAP2*. Somatic mutations were annotated using ANNOVAR, and an APG was labelled mutated if it carried any non-synonymous, frameshift, stop-gain, or stop-loss variant within coding exons. Gene-level copy number was inferred from Battenberg output. Ultimately, a sample was classified as immune-escaped if it exhibited at least one of: HLA mutation, HLA LOH or APG mutation. HLA AI was not considered indicative of immune escape, as AI can result from subclonal LOH or focal gains, making its impact on antigen presentation uncertain. For samples in which HLA alterations could not be fully evaluated, but neither HLA nor APG alteration was detected, the immune escape status was recorded as unknown, reflecting that a possible immune escape genotype could not be ruled out.

Neoantigens were predicted using the Python-based pipeline NeoPredPipe^89^, which integrates ANNOVAR^90^ and netMHCpan (v4.0)^91^. In summary, all somatic SNVs and indels were first annotated with ANNOVAR, and mutated peptide sequences were generated for all non-synonymous exonic mutations. For each mutation, we considered all 9- and 10-mer peptides spanning the altered amino acid(s), resulting in either (i.) a 19-amino acid window for SNVs or (ii.) the peptide extending to the next predicted stop codon for frameshift mutations. These peptides were assessed for both novelty and predicted binding affinity to the patient-specific six-allele HLA genotype (comprising HLA-A, HLA-B, and HLA-C genes). Peptides that were novel relative to the healthy human proteome and had a binding rank ≤2 (within the top 2% of binders compared with random peptides) were designated as neoantigens. All patient HLA alleles were included in predictions, irrespective of mutation or LOH status at the HLA locus. 29 samples were excluded from neoantigen calling because netMHCpan could not predict at least one of their HLA haplotypes. A mutation was classified as a neoantigen if at least one of its downstream mutated peptides was predicted as a neoantigen for any of the patient’s six HLA alleles. Neoantigen burden was defined as the total number of neoantigenic mutations per sample. We assessed the immunogenicity of individual mutations in each patient using the patient harmonic-mean best rank (PHBR) score metric, which accounts for all novel peptides generated by a mutation as well as all HLA alleles present in the sample. Low PHBR values indicate mutations that are more likely to be presented on the cell surface and therefore have higher immunogenic potential, whereas high PHBR values indicate lower immunogenicity. The overall immunogenic potential of a mutation within a cohort is defined as the median PHBR value across all samples in that cohort. For each mutation-HLA haplotype pair, we generated all overlapping 8–11-mer peptides spanning the mutation and evaluated their binding affinity to the HLA allele using the ‘all-predictions’ mode of NeoPredPipe, recording the best (lowest) rank for each. For each patient, PHBR values were calculated as the harmonic mean of the six best rank values corresponding to their six HLA haplotypes, counting homozygous alleles twice. PHBR values were computed for all single-nucleotide mutations located in driver genes observed in at least four cancers in the cohort. Samples with incompatible HLA alleles (n= 29) were excluded from analysis. To investigate the impact of HLA alterations on PHBR values, we repeated this analysis in samples affected by such alterations, using only the subset of incomplete (<6) HLA allele profiles. To assess patient- and HLA-dependent selection on driver genes, we compared PHBR values for mutations in these genes between patients carrying the mutation and those who did not. Negative differences indicate that mutations preferentially occur in patients for whom they have lower immunogenic potential. Comparisons between PHBR values in mutated and non-mutated patients were performed using the Wilcoxon rank-sum test, with Benjamini–Hochberg correction applied for multiple testing. For analyses of PHBR within subgroups, the immunogenic potential of individual mutations was quantified using the median PHBR values across all patients. PHBR values of the same mutation across different cohorts were compared using paired Wilcoxon rank tests.

Because the distinct EC molecular subtypes (CN-low, CN-high, POLE and MSI) exhibit markedly different mutational and immune profiles, statistical analysis on PHBR scores was performed after stratification by subtype. Pairwise comparisons were conducted using Wilcoxon tests. Multiple regression was conducted to assess the relationship between immune escape types and neoantigen burden. These used the lm function on the logarithm of neoantigen burden, allowing estimation of the fold change in burden associated with each escape type. Similar regression analyses that included clinical covariables were performed using the logarithm of total mutation burden as an additional independent variable.

### Microbial identification

Unmapped reads from the BAM files were analyzed using Centrifuge (version 1.0.4)^92^ with the h+p+v+c genome index, which consists of human, prokaryotic, and viral genomes (including SARS-CoV-2), downloaded from https://zenodo.org/records/3732127. Two non-default parameters were applied: ‘--host-taxids 9606’ to specify the human taxonomy ID, ensuring that any reads aligning to the human genome were assigned exclusively to Homo sapiens; ‘--min-hitlen 35’ to increase the minimum number of aligned bases from the default value of 22 to 35. Centrifuge outputs were converted into Kraken-style summary reports using the ‘centrifuge-kreport’ utility. The total number of human reads per sample was computed as the sum of (i) reads mapped to the human genome in the original BAM files and (ii) unmapped reads subsequently classified as human by Centrifuge. Species- and genus-level read counts were then extracted from the reports and normalized to one billion total human reads per sample. Species-level quantification data were further processed to assess associations between microbial composition and clinical as well as technical covariates. To this end, a permutational multivariate analysis of variance (PERMANOVA) was performed to determine the extent to which variation in microbiome structure could be explained by the following predictors: (i.) molecular subtype (TCGA), (ii.) tumor stage, (iii.) tumor grade, (iv.) histotype, (v.) sequencing facility. Most of the variance could be explained by the sequencing facility (pseudo-F = 10.8, p < 0.001), followed by histotype (pseudo-F = 1.5, p = 0.002). As we suspected possible contamination to be the main source of microbial abundance in biopsy samples, additional UMAP clustering was performed to reveal further dependency on the sequencing facility. The formation of local clusters of samples annotated by sequencing facilities suggests the presence of batch effects, potentially driven by facility-specific microbial contamination or other technical artefacts. To assess the contribution of the mentioned sources clinical and technical variables to variation in microbial abundance, variance partitioning was performed using mixed linear models as implemented in the variancePartition R package (v1.32.5). The proportion of variance explained by each variable was computed for every species, decomposing the total variability into biological and technical sources. Most species showed residual variance exceeding 75%, indicating a substantial proportion of individual variability that was not explained by the included variables. Among the tested features, the sequencing facility accounted for the largest share of explained variance, consistent with the results obtained from PERMANOVA.

### Mitochondria Analysis

Mutect2 v4.1.0.0 (GATK v4.5.0.0)^93^, with default settings was used to call somatic mitochondria DNA (mtDNA) SNVs and indels. mtDNA variants were excluded if they met any of the following criteria: 1) had low mapping or base quality score (<20); 2) An alternative allele frequency <1%; 3) Missing alternative reads in any strand direction, or 4) were located in hypermutated regions; 302-316, 514-525 or 3106-3109.

Where purity and ploidy could be estimated, the mitochondrial DNA copy number was estimated as per Yuan et al.^48^:

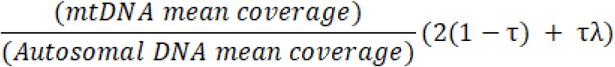

given sample purity (τ), tumour ploidy (λ), and autosomal and mitochondria mean coverage depth estimated by fastMitoCalc^94^. Mutational selection of mitochondrial protein-coding genes was evaluated via the non-synonymous to synonymous mutation rate (dN/dS) using dNdScv^18^. The mitochondrial genome was isolated as the reference genome and default parameters were used. As the *MT-ND6* gene was subjected to replication bias, it was excluded when calculating the global mitochondrial dN/dS.

### Unsupervised clustering

Clustering was performed to identify similar EC subgroups based on integrated genomic and clinical data. The clustering input comprised molecular signature burden proportions for SBS, DBS, ID, CN and SV; SSV burden (SNVs and indels); percentage genome altered; number of structural variants; age; genome-wide LOH fraction; ploidy and purity; and driver genes. To limit model complexity and reduce the risk of overfitting, we selected a subset of drivers using an unsupervised feature selection approach based on the Laplacian Score via the Python GitHub repository LaplacianScore^95^. Specifically, Laplacian Score was applied to the set of 107 EC driver genes to identify features that best preserve the intrinsic similarity structure of the data. Based on these gene features, pairwise similarities between all samples were calculated using euclidean distance, and a k-nearest neighbor graph (k = 5) was subsequently constructed across all samples, following the default parameter settings of the implementation. This graph models the local manifold structure of the data, which is then used to compute the Laplacian Score for each feature. Features that better respect the intrinsic structure show lower Laplacian score values and are therefore considered more informative for downstream analysis. The resulting ten most informative, EC-associated genes were found to be, in descending order, *PTEN*, *TP53*, *PIK3CA*, *ARID1A*, *POLE*, *SYNE1*, *PIK3R1*, *DYNCH1*, *AKAP9*. For these genes, we included all nonsynonymous variants for the genes annotated as oncogenic by OncoKB and collapsed them into a binary mutational status. Prior to clustering, all variables were scaled to a range between 0 and 1. Hierarchical clustering was performed on these features using the pvclust R package (v2.2.0), evaluating all combinations of the following agglomerative methods and distance measures: (i.) Agglomerative Method: ward.D and ward.D2 and (ii.) distance measure: euclidean, correlation (1-Pearson correlation) and canberra. The combination of ward.D2 linkage with canberra distance recapitulated known TCGA molecular subtypes best and revealed meaningful subclusters for TCGA subtypes MSI and CN-high. This configuration was therefore selected for hierarchical clustering.

Clustering results were visualized using a dendrogram and per-sample feature tracks generated with the R package complexHeatmap (v2.18.0), with samples ordered according to the hierarchical structure. The resulting dendrogram was cut to obtain seven clusters. The choice of number of clusters was guided by both silhouette scores and biological interpretability. Major splits at higher dendrogram heights (splits 1–3) corresponded to the four established TCGA molecular subtypes CN-high, CN-low, MSI, and POLE, mainly reflecting large-scale differences in genomic instability and mutational burden. Further splits allowed to capture novel subgroups of CN-high (WGD), CN-high (focal LOH), HRD and MSI-*POLD1* subtypes. Cluster stability for these subgroups was confirmed by silhouette widths, which showed positive means for all subgroups.

### Survival analysis

To visualize survival differences across TCGA and novel sample subgroups, Kaplan-Meier (KM) curves were generated using ggsurvplot from the ggsurvminer package. Log-rank tests were applied to compare survival between groups. KM plots included risk tables and p-value annotations to facilitate interpretation. Associations between age molecular features (including molecular subgroup, mutational burden, genomic instability and EC driver genes) and survival outcomes were evaluated using Cox proportional hazards (PH) regression models. Both univariable and multivariable analyses were performed to adjust for confounding biases. Multivariable EC driver models were adjusted for covariables age, EC molecular subtype, SNV burden and PGA as indicated. Hazard ratios (HRs) with 95% confidence intervals and two-sided *P* values were reported. Proportionality of hazards assumption in Cox models was confirmed by scaled Schoenfeld residuals. Statistical significance was defined as *P*<0.05.

### Actionability of driver gene mutations and networks

The OncoKB^54^ and COSMIC Mutation Actionability in Precision Oncology Product^55^ databases were queried to evaluate the therapeutic implications of genetic events. These databases catalogue approved marketed drugs, which have shown efficacy in the context of specific driver gene mutations, based on clinical trial and other published evidence. OncoKB also provides biological evidence supporting the cancer driver gene as being predictive of a response to a given drug. To perform a chemogenic analysis of cancer networks for each cancer type, we used protein products of cancer driver genes was used to seed a search for interacting proteins in the canSAR interactome, as previously described^56^. Essential and selective genes, including those with lineage specificity, were identified using the ShinyDepMap analysis server. To identify candidate synthetic lethality relationships for tumour suppressor drivers, Synthetic Lethal Vulnerabilities Exhibiting Reciprocation (SLYVER)^57^, was used which applies pan-cancer tumour cells lines in the DepMap CRISPR-Cas9 screen data that mirror alterations in the TCGA uterine corpus endometrial carcinoma cohort.

### Functional analysis of *INNPL1* (SHIP2) mutations

Human endometrial adenocarcinoma cell lines HEC-1A, EN, KLE and SNG-II cells were a kind gift from Konstantin Dedes (University of Zurich, Switzerland). HEC-1B, Ishikawa, MFE-296 and NOU1 cells were shared by Britta Weigelt (previously Cancer Research UK, Lincoln’s Inn Fields UK). MFE-280 cells were purchased from the European Collection of Authenticated Cell Cultures (Porton Down, UK). HEC-59 cells were obtained from the JCRB cell bank (Osaka, Japan). hEM3 cells were a kind gift from Tian-Li Wang (Johns Hopkins University School of Medicine, USA).

All cell lines were culture in an atmosphere of 5% CO_2_ at 37 °C. SNG-II were cultured in DMEM medium with 10% foetal bovine serum (FBS); EN in DMEM with 20% FBS; MFE-280 and MFE-296 in 40% RPMI/40% DMEM with 20% FBS; KLE and Ishikawa in DMEM/F12 with 10% FBS; HEC-1-A and HEC-1-B in MEM with 10% FBS; NOU1, and HEC-59 in MEM with 15% FBS. Cell line identity was confirmed by STR profiling. The MSI status was determined using the Promega MSI analysis system and *INPPL1* mutation by Sanger sequencing. siRNAs targeting GAPDH, INPPL1/SHIP2 and non-targeting control were purchased from Dharmacon. Cells for immunoblotting were washed with ice cold PBS and then lysed in RIPA buffer with protease and phosphatase inhibitors (Pierce Halt Cocktail, ThermoFisher, Waltham, MA, USA). Lysate protein concentration was determined using the CB-X. Protein Assay (GBiosciences, Saint Louis, MO, USA). Proteins (30 mg) from each sample were separated using NuPAGE 3-to-8%, Tris-Acetate gels (Invitrogen, Waltham, MA, USA) and blotted onto PVDF membranes (Millipore, Burlington, MA, USA) by wet transfer. Membranes were blocked with 5% dried milk in 0.1% TBST for 1 h, incubated with primary antibody overnight at 4 °C washed with TBST x 3, incubated with actin loading control antibody for 1 h at room temperature, washed x 3 and incubated with secondary antibody in 5% dried milk at room temperature for 1 h. Membranes were then imaged using the LI-COR imaging system (LI-COR Bioscience, Lincoln, NE, USA). Antibodies used were primary: anti-human SHIP2 (C76A7) (2839, Cell Signaling Technology (Beverly, MA, USA), 1:1000), anti-human PTEN (9559, Cell Signaling Technology, 1:1000), anti-AKT (9272, Cell Signaling Technology, 1:1000), anti-pAKT^S473^ (4060, Cell Signaling Technology, 1:1000), anti-ERK1/2 (9102, Cell Signaling Technology, 1:1000), anti-pERK1/2^T202/T204^ (4370, Cell Signaling Technology, 1:1000), anti-human GAPDH antibody (5174, Cell Signaling Technology, 1:1000), anti-human Actin antibody (ab8227, AbCam, Cambridge, UK, 1:5000) and secondary: Goat Anti-Rabbit IgG (LIC-925-68071, LI-COR Bioscience, 1:15000). Densitometry was performed using ImageJ (NIH).

### Data availability

Data from the National Genomic Research Library (NGRL) used in this research are available within the secure Genomics England Research Environment. Access to NGRL data is restricted to adhere to consent requirements and protect participant privacy. Data used in this research include:

- BAM files and VCF files containing SNV, indel and SV calls. The corresponding metadata and file locations for these files can be obtained through LabKey by querying the ‘cancer_analysis’ table.
- Processed clinical and genomic data, available in the Research Environment within the folder /published_data_archive/paper_data/paper_data_RR27

To support reproducibility, a README file listing all participants’ IDs used is included in /published_data_archive/paper_data/paper_data_RR27

At present, there is no proposed end date for data access within the research environment.

Access to NGRL data is provided to approved researchers who are members of the Genomics England Research Network, subject to institutional access agreements and research project approval under participant-led governance.

The raw data, including patient profiles and corresponding genomic sequencing data, are available under restricted access for patient privacy reasons. Access can be obtained by first applying to become a member of the Genomics England Research Network (https://www.genomicsengland.co.uk/research). The process for joining the network is described at https://www.genomicsengland.co.uk/join-us. Your institution will need to sign a participation agreement available at https://files.genomicsengland.co.uk/documents/Genomics-England-GeCIP-Participation-Agreement-v2.0.pdf and email the signed version to. Once you have confirmed your institution is registered and have found a domain of interest, you can apply through the online form at https://www.genomicsengland.co.uk/join-us, where you can specify the reason for access and expected time frame that you wish to have access. Once your Research Portal account is created, you will be able to log in and track your application. Your application will be reviewed within 10 working days. Your institution will validate your affiliation. You will need to complete the online Information Governance training and will be granted access to the Research Environment within 2 days of passing the online training. The Research Environment is accessed through Amazon WorkSpaces (https://clients.amazonworkspaces.com/). For more information on data access, visit https://www.genomicsengland.co.uk/research.

